# ‘Genome delivery of a contractile tailed phage and its superinfection exclusion mechanism’

**DOI:** 10.64898/2026.03.06.709160

**Authors:** Aritz Roa-Eguiara, Leyre Marín-Arraiza, Victor Klein-Sousa, Mònica Santiveri, Nicole R. Rutbeek, Damien Piel, Tillmann Pape, Nicholas Sofos, Ivo Alexander Hendriks, Michael Lund Nielsen, Haidai Hu, Alexander Harms, Nicholas M. I. Taylor

## Abstract

Successful viral infection requires efficient adsorption to the target cell, followed by membrane penetration for genome translocation into the host cytoplasm. Bacteriophage T4 initiates infection by binding to Escherichia coli receptors, triggering sequential conformational changes that culminate in genome delivery through viral channels spanning the host cell envelope. Here, we resolved structures of bacteriophage T4 at discrete stages. This revealed how long tail fiber extension affects the baseplate to initiate tail contraction, and the channel formation by the tape measure protein for genome translocation. We further demonstrate that the virus-encoded superinfection exclusion protein Imm binds to the tape measure protein to prevent secondary infections. Our findings offer insights into the coordinated process of genome delivery and phage-encoded superinfection exclusion proteins that prevent genome translocation.

## Introduction

A successful viral infection depends on the ability of the virus to recognise and penetrate the host cell membrane to deliver its genome and replicate. Bacteriophages (or simply phages) are viruses that infect bacteria and have evolved a variety of mechanisms to achieve this^1,2^. Phages with long tails are the most abundant morphotype known to date and are further classified into myoviruses and siphoviruses, corresponding to contractile and non-contractile tailed phages, respectively^3^. Their distal tail ends assemble receptor binding proteins (RBPs) used to mediate recognition and attachment to their host. Following adhesion, the tail complex undergoes conformational changes to inject the viral genome into the host cytoplasm^4^. These phages encode a tape measure protein (TMP) which acts as a structural scaffold defining tail length^5^. Beyond its structural function during phage assembly, a role in membrane fusion and genome translocation has been proposed, although direct observation has remained elusive^6–10^.

Phage T4 is a model myovirus infecting *Escherichia coli K-12* and *B* strains^11^ that has been extensively studied in fundamental biology^12,13^. To initiate infection, phage T4 uses its long tail fibres (LTFs)^14–17^, a set of RBPs, for reversible binding to the outer membrane protein C (OmpC) and/or lipopolysaccharides (LPS)^11^. Binding of LTFs triggers the rearrangement of the baseplate, located at the distal end of the tail, to irreversibly attach to the cell surface through the short tail fibres (STF)^18–20^, a second set of RBPs that bind LPS. The transition from a high- to a low-energy state of the baseplate initiates sheath contraction, driving the inner tube through the bacterial envelope and forming a conduit for genome translocation^21–25^. The precise signal transduction from receptor binding by the LTF to tail sheath contraction remains uncertain despite receiving considerable attention^18,26–29^. Afterwards, it has been observed that the tube binds the inner membrane, followed by genome translocation. However, the detailed mechanism of membrane penetration is not known^8,10,24^.

These steps summarise the initial early infection process, which can be targeted by some superinfection exclusion proteins that prevent secondary infections of the same cell. This is the case for the immunity protein (Imm) of phage T4, among others^30–32^. The Imm is a small, 9.3 kDa membrane protein produced by phage T4 and has been shown to block genome translocation^33,34^. However, the mechanism through which this occurs remained to be described.

We hypothesised that the TMP of phage T4, also named gp29, regulates genome translocation through pore formation in the inner membrane. Furthermore, we theorised that the Imm functions by interfering with this process^35^, performing a similar role to immunity proteins of pore forming colicins^36–38^.

In this research, we use cryo-electron microscopy (cryo-EM) and structural predictions to study the intact structures of phage T4 during early infection. We elucidate the conformational dynamics of the LTFs in the context of baseplate rearrangement. Additionally, we resolve the structures of two post-contracted states, before and after genome ejection, revealing the basis of inner membrane recognition by the TMP and the regulation of genome translocation. Furthermore, we provide structural and functional insight into how the Imm prevents secondary infections by directly binding to the TMP. Together, and building on a large collection of previous research, this allows us to construct a molecular movie for the molecular choreography that allows T4 to breach the cell envelope and inject its genome into the host cell.

## Results

### Structures of phage T4 at different steps of DNA ejection

Phage T4 undergoes a plethora of conformational rearrangements during initial infection and regulates genome delivery into the host^18,19,24,39^. To study this, we characterised three different phage T4 samples that represent different states during infection initiation: the pre-contracted virion with retracted LTFs, the post-contracted virion before genome ejection, and the post-ejected genome state of the post-contracted virion upon lipid membrane interaction. The phage sample was analysed by mass spectrometry, confirming that all the modelled proteins were present in the purified samples (**Supplementary Table 1**).

First, we purified the phage particles to study the fully assembled virion on its extended state by local reconstruction of phage subcomplexes using single particle cryo-EM (**Extended Data Fig. 1,2, Supplementary Table 2**). Next, we triggered sheath contraction via urea treatment and reconstructed this state prior to DNA ejection. Finally, the post-ejected genome state was obtained by mixing the contracted virion with *E. coli* lipids harboured in nanodiscs (**Extended Data Fig. 3,4, Supplementary Table 2**).

Initially, we determined the structure of the pre-contracted virions, which have the LTFs retracted towards the phage tail, and their structure provides insights into the interaction of LTFs with the rest of the virion (**Fig. 1a**). We modelled the tail-neck-portal assembly (**Extended Data Fig. 5a**), and analysed the structure of gp13 in the mature virion, for which the domain organisation is similar to gp11 of phage SU10^40^ and gp81 of podovirus GP4^41^ (**Extended Data Fig. 5b**). We resolved the binding of gp13 to the fibritins. This interaction occurs via a flexible loop of gp13 (fibritin dock loop; spanning gp13 residues 196-240)that adapts to the conformation of the collar, the fibritins that bend around the neck, and the whiskers which bend towards the tail end^15,22^ (**Extended Data Fig. 5c**). Analysis of the neck-capsid binding revealed two assemblies, differing from each other by a ∼4° rotation between the neck-portal and the capsid (**Extended Data Fig. 5d,e**). Furthermore, the structure of the fully assembled virion reveals novel interactions between the tube and the baseplate-sheath interface, anchoring the tube to the hexagonal baseplate and pre-contracted sheath (**Extended Data Fig. 5f**).

**Figure 1.**
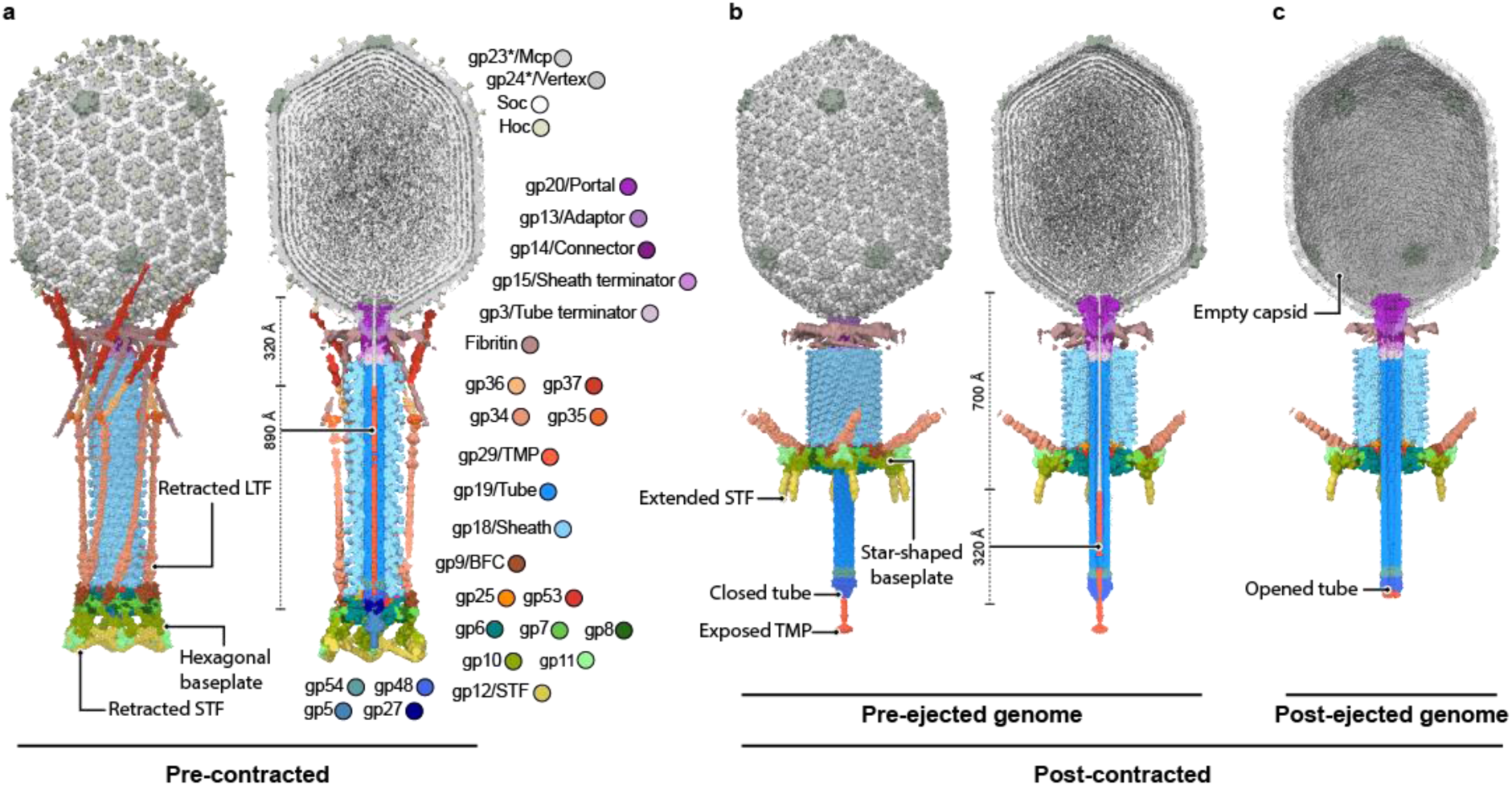
Phage T4 states and genome release. **a**, Pre-contracted conformation of the phage with the LTF in the retracted state. On the left side of the panel, is shown the surface representation of the atomic models fitted in the low-resolution reconstruction of the tail, neck and portal. Capsid and N-terminus of gp37 are shown as electrostatic potential maps. On the right, sliced maps of the tail and the capsid. (Mcp, major capsid protein; TMP, tape measure protein; BFC, baseplate-fibre adaptor protein; STF, short tail fibre) **b**, Post-contracted state of the virion, displayed in the same manner as in panel a. **c,** Sliced electrostatic potential maps of the genome ejected virion after supplementation with nanodiscs harbouring *E. coli* lipids.

We then determined the structure of the post-contracted virion before genome ejection (**Fig. 1b**). The conformational changes upon phage tail sheath contraction release part of the genome from the capsid into the tube. However, the DNA is retained inside the virion, as shown previously^24^ (**Extended Data Fig. 6a,b**) During this process, the portal extends its crown helices, displacing the portal loop from the lumen (**Extended Data Fig. 6c**). Additionally, we observed that the TMP is partially exposed to the medium (**Extended Data Fig. 6d,e**).

Subsequently, we triggered genome ejection by supplementing the contracted sample with *E. coli* lipid membranes^23,42^ (**Fig. 1c**). The interaction with the lipids releases the tube content together with part of the genome which leaves the capsid through the tube (**Extended Data Fig. 6f**). Some fraction of the genome remains wrapped around the portal (**Extended Data Fig. 6g**) and interacts with the internal protein III (Ip3), an anti-phage defence protein^43^ (**Extended Data Fig. 6h,i**). We also observed an unknown density above the portal, previously described in *in situ* infection^24^ (**Extended Data Fig. 6j**). This indicates that membrane recognition alone is not sufficient for the complete injection of the genome into the host cell^2^ and that Ip3 is probably injected after all of the genome has been translocated.

Together, these structures describe the rearrangements of the fully assembled virion of phage T4 along genome translocation, suggesting a regulated release of the phage genome upon membrane recognition. We further dived into the molecular details of each state and their relevance in infection initiation.

### Long tail fibre extension is coupled to baseplate conformational changes

To study the interactions between the LTF and the rest of the virion, we first characterised their retracted state towards the tail. Fibres are often flexible proteins^43^, a characteristic that hampered a single local reconstruction of the entire LTF (**Fig. 2a**). To overcome this, we performed local refinements along its structure to improve the quality of the maps (**Extended Data Fig. 7a,b**). This led us to resolve the overall structure of the kneecap protein (KC), gp35, and the LTF domains.

**Figure 2.**
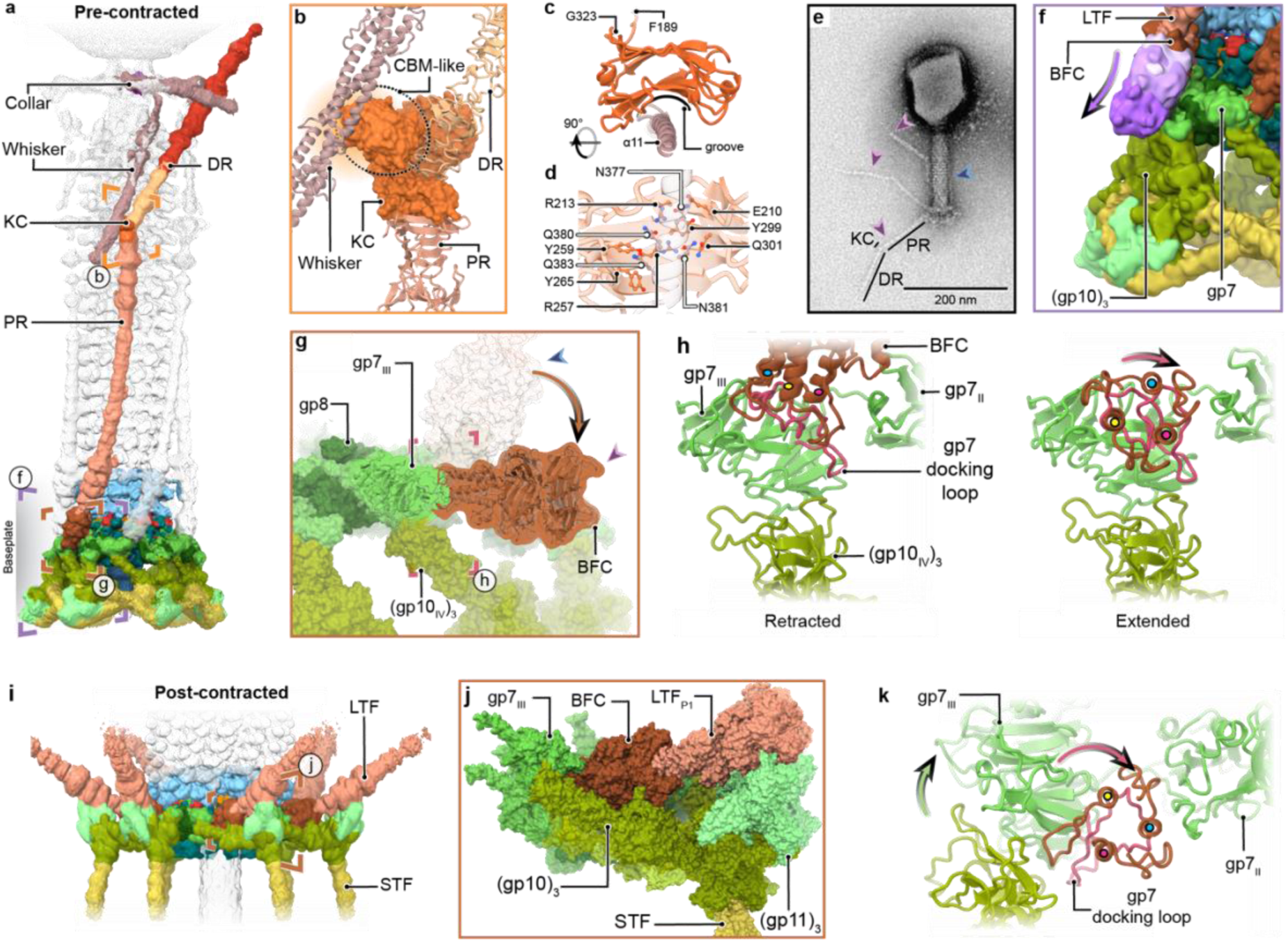
Assembly and extension of the LTF. **a**, Cryo-EM map of the focus refinement of a complete LTF, low-pass filtered to 8 Å and low threshold (σ5) (DR: Distal Rod of the LTF, PR: Proximal Rod of the LTF, KC: Kneecap protein). **b**, KC interacting with the whisker, distal, and proximal rods of the LTF. **c**, Whisker helix 11 (α11) interacting with the groove of the CBM-like domain. **d**, Front view of the groove in panel b with the side chains from polar and charged residues of the interface for referenced but not resolved in the ∼7 Å resolution map. **e**, Negative stained phage T4 sample imaged by TEM at high salt concentration, showing extended and retracted fibres (purple and blue arrows respectively). **f**, Cryo-EM maps from the BFC classes, filtered to 10 Å and σ3 for comparison, from the retracted LTF (brown) to the extended BFC state (gradient of purple from light to dark). **g**, Comparison between the retracted and extended state of the BFC. **h**, Focus view of the gp7 docking loop binding to the retracted and extended BFC (*left* and *right* respectively). Coloured circles on top of BFC N-terminal helices to assist tracking movement between states. **i**, Cryo-EM map of the post-contracted phage T4 tail focused on the star-shaped baseplate, low-pass filtered to 10 Å and low threshold (σ5). **j**, Side view of the baseplate wedge in surface representation to show the interaction between the LTF_P1_ and the peripheral baseplate. **k**, Focus view of the gp7 docking loop on the star-shaped baseplate, as in panel h.

The KC joins the proximal rod, formed by (gp34)_3_; and the distal rod, formed by (gp36)_3_-(gp37)_3_ (**Fig. 2b**). The KC folds into three domains: i) a tower-like domain similar to the fold of the C-terminal region of the proximal rod (tower-like domain; KC (1-21 and 324-372)), ii) an ILEI/PANDER-like domain (KC (22-188)) similar to eukaryotic signalling molecules (InterPro: IPR039477)^44–46^, and iii) a carbohydrate binding module-like domain (CBM-like; KC (189-323)) that binds the whisker α11 (fibritin (361-386)) (**Fig. 2c, Extended Data Fig. 7c,d**). The CBM-like domain forms a structurally conserved groove among CBMs, where carbohydrate ligands bind in other studied CBMs^47,48^ (**Extended Data Fig. 7e**). The groove exposes a series of polar and charged residues (E210, R213, R257, Y259, Y265, Y299, Q301) facing the polar side chains of one whisker subunit (N377, Q380, N381, Q383) (**Fig. 2d**). This data shows the role of the KC in preserving LTF assembly and suggests its role in the stabilisation of the retracted conformation through electrostatic interactions.

To study the infection process in detail, we next investigated the extension of the LTF from the tail by negative stain transmission electron microscopy (TEM). It has been previously observed that the extended conformation of the LTF is favoured under high-ionic strength^15^. Therefore, we increased the salt concentration of the vitrification buffer from 8 mM to 350 mM MgCl_2_ via dialysis, inducing extension of the LTFs (**Fig. 2e**). We reversed the salt concentration by dialysis of the samples to 8 mM MgCl_2_, a condition that led to retraction of the LTFs (**Extended Data Fig. 7f,g**). This data confirms a dynamic LTF response to environmental changes and sensitivity to ions that can regulate the exposure of the fibres.

The range and path of extension of the LTF is dictated by its attachment to the baseplate. The LTF binds to the baseplate protein gp7 through the baseplate-fibre connector (BFC), (gp9)_3_. More precisely, the BFC is binding to domain III of gp7 (gp7_III_; gp7 (172-537) via the docking loop of this domain (gp7 (258-292)). First, we observed that in our baseplate reconstruction, 14% of the BFCs were not in the retracted conformation. We classified the position of the BFC at each wedge and refined three discrete orientations (**Fig. 2f,g**). Finally, analysis of the individual conformations indicated that the extension of the LTF follows a defined tilting angle coupled to a rotation of the fibre (**Fig. 2h**). Therefore, this movement extends the fibre away from the tail and orients the receptor binding domain of the LTF to align the tail perpendicular to the membrane.

To better understand the complete range of motion of the fibres, we next studied the BFC-gp7_III_ interaction in the post-contracted baseplate (**Fig. 2i,j**). The movement of the docking loop during LTF extension culminates in a turn of ∼90° in the star-shaped baseplate, compared to its extended position in the hexagonal baseplate (**Fig. 2k**). This rotation of the docking loop would be first transmitted to gp7_III_ by turning and lifting gp7_III_ from the baseplate plane (**Extended Data Fig. 7h**). At the same time, gp7_III_ is directly binding to the domain IV of (gp10)_3_ (gp10 (406-602)) that connects with the peripheral baseplate. Overall, analysis of the hexagonal and the star-shaped baseplate conformations reflect how the extension movement of the LTF-BFC likely exerts a pulling motion from gp7_III_ to initially transmit the signal to the peripheral baseplate, leading to the intermediate state previously reported by cryo-electron tomography^24^.

### The tube is anchored to the hexagonal baseplate

To further understand the initiation of sheath contraction, we compared the baseplate–sheath interface in pre- and post-contracted virion states. In phage T4, the N-terminal anchor loop of the tube initiator protein gp48 is an integral component of this interface in the pre-contracted conformation (**Fig. 3a**). Specifically, the anchor loop of gp48 (gp48 (7-32)) inserts between the platform domain of gp53 (gp53 (89-168)) and the baseplate-proximal layer of the sheath (**Fig. 3b,c**), thereby restricting conformational changes. Following irreversible binding of the STFs to host LPS^19,20^, the baseplate undergoes axial expansion (**Fig. 3d,e**), which disengages the tube from the baseplate^18^. This rearrangement vacates the interface previously occupied by the gp48 anchor loop, enabling sheath protomers to interact with gp53 and initiate contraction (**Fig. 3f**). Essentially, the gp48 anchor loop helps to secure the distal end of the tube within the hexagonal-baseplate architecture until its release upon baseplate rearrangement, which permits sheath contraction and subsequent membrane penetration.

**Figure 3.**
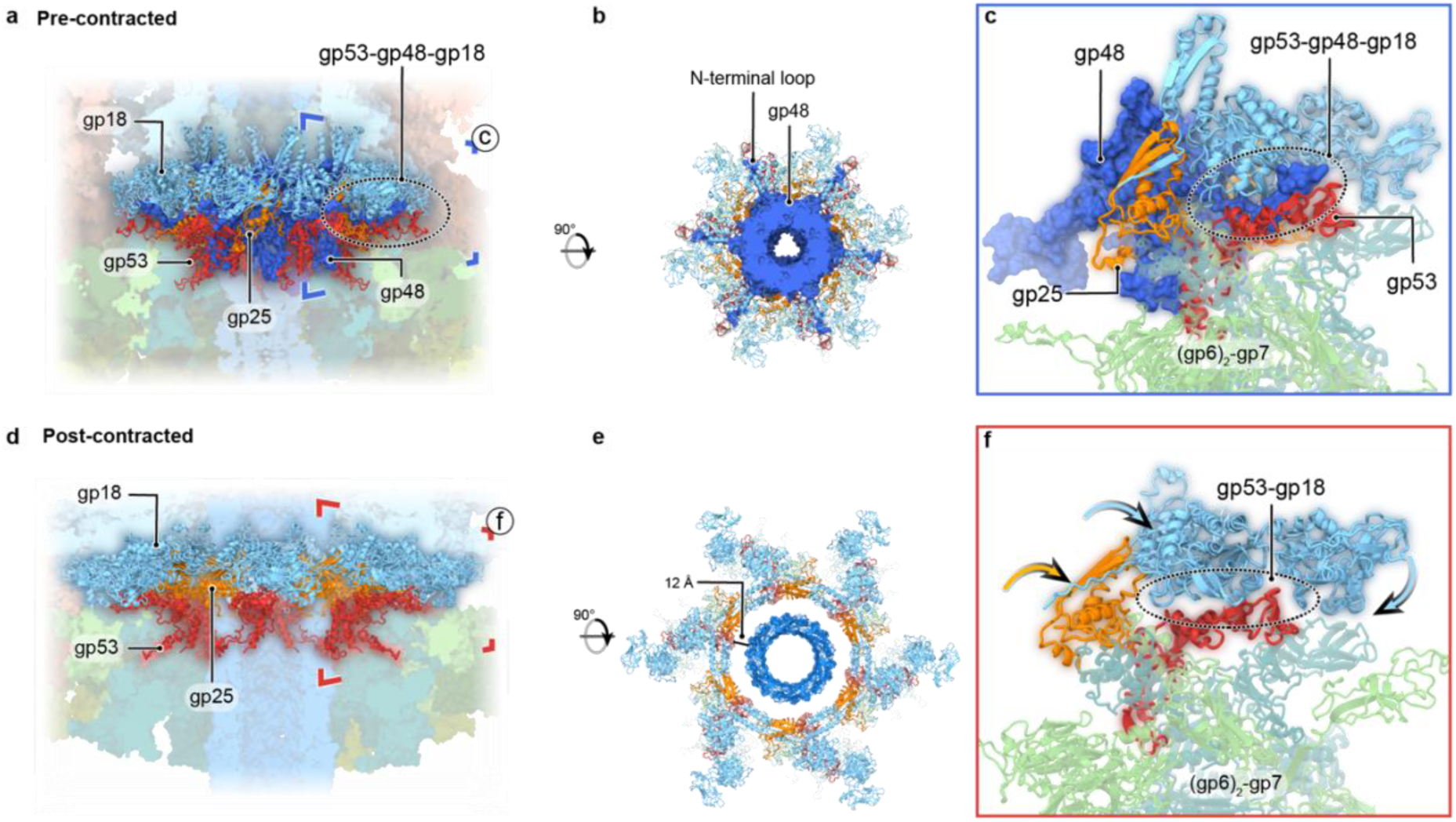
Inner baseplate, sheath and distal tube end disengagement. **a**, Organisation of the baseplate-sheath interface. Gp48 is shown in surface representation. **b**, Top view of the gp48 section showing the N-terminal insertion loop into the inner baseplate. **c**, Isolated view of the N-terminal gp48 insertion into the inner baseplate (encircled). **d-f**, Same view of the inner baseplate as in panels a-c, but of the post-contracted virion state.

### The distal end tube and the TMP regulate genome release

To study the translocation of the genome, we next examined how the virion prevents spontaneous DNA ejection immediately after contraction. We combined the information from the virion reconstruction, the sequence of the TMP and AlphaFold2 predictions to estimate the location of the TMP domains inside the tube at each stage. This allowed us to identify an N-terminal coiled-coil helical domain (TMP_CC_; TMP (1-120)) closest to the genome terminus, a putative transmembrane segment (TMP_TM_, TMP (207-246)) supported by transmembrane prediction, a helical bundle domain (TMP_HBD_; TMP (273-518)) and a C-terminal loop (TMP_CL_; TMP (571-590)) (**Extended Data Fig. 8a-c**).

Initially, the pre-contracted virion harbours six copies of TMP_CL_ in two distinct conformations. Three of them are bound to the distal end of the tube, while the other three fragments are only partially visible and arrange in a different conformation (**Fig. 4a**), similar to what has been observed for phage HFTV1^49^. These are the only fragments that could be built *de novo* in the pre-contracted virion.

**Figure 4.**
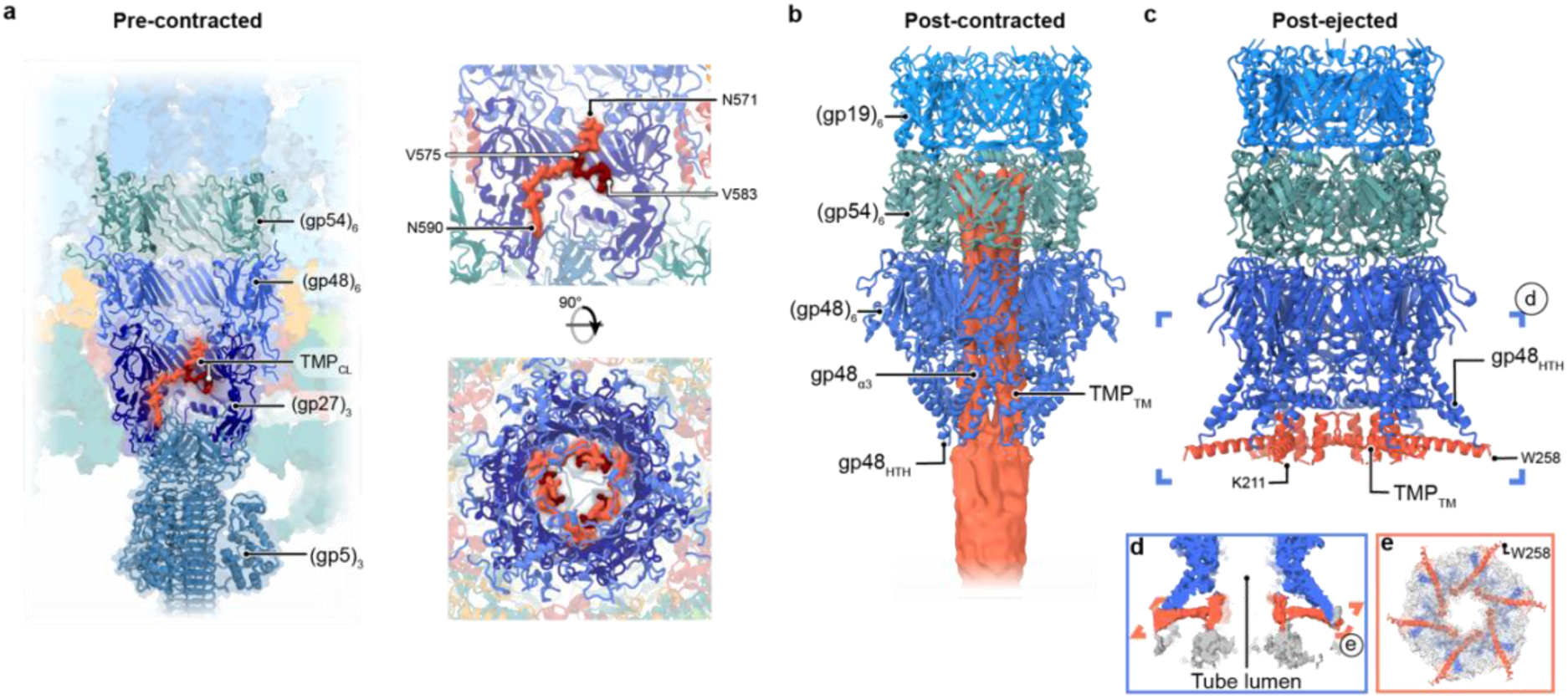
Genome ejection regulation by the TMP. **a**, Organisation of the baseplate–hub (gp5_3_-gp27_3_-gp48_6_-gp54_6_) (*left*), and detailed view of the densities assigned to the TMP_CL_, from side and top (*right*). **b**, Genome retention mechanism by the TMP, most likely the TMP_TM_, binding to the distal end tube, (gp48)_6_, on the post-contracted virion. **c**, post-ejected genome conformation of the distal end tube and the TMP_TM_ helix bound to gp48. **d**, Cryo-EM map coloured as in panel c, showing the densities for the interface between the tube and the TMP. The unresolved density in grey, showing the formation of a conduit. **e**, Bottom view of the tube opening of the cryo-EM map at low threshold to show the densities observed around the model.

In the contracted sample, the baseplate hub subcomplex gp5.4-(gp5)_3_-(gp27)_3_, which forms the puncturing device, is released and the TMP_HBD_ is exposed to the outside of the tube lumen (**Fig. 4b**). In this conformation, the distal end tail tube protein gp48 binds to the TMP arresting the ejection process. This is achieved by constraining the tube lumen with the helix α3 of gp48 (gp48_α3_; gp48 (170-178)) and closing its helix-turn-helix fold (gp48_HTH_; gp48 (241-282)) against a TMP helical region, presumably the TMP_TM_ (**Extended Data Fig. 8d**). The previously described displacement of the stopper loop in the portal (**Extended Data Fig. 6c**) emphasizes the role of the (gp48)_6_-TMP_6_ in retaining the DNA inside the virion, as it is the only observed constriction point. In conclusion, these interactions between the TMP_TM_ and the distal end tail tube show how the contracted virion arrests genome ejection after contraction.

We then aimed to determine how the post-contracted phage interacts with lipid bilayers, resembling the final injection state. When we incubated contracted phage T4 with nanodiscs containing *E. coli* lipids, we observed one conformation forming an open conduit for genome release (**Fig. 4c**) and two other alternative states (**Extended Data Fig. 8e,f**). In the open state, (gp48)_6_ acquires a flared conformation upon membrane binding, where each gp48_HTH_ is expanded outwards and interacts with part of the TMP. We could assign the TMP density to the TMP_TM_, as supported by bioinformatic structural and transmembrane prediction (**Extended Data Fig. 8g,h**). This reinforces the previous observation of the post-contracted state before ejection, the TMP_TM_ region remains interacting with the distal end tail tube after contraction and throughout the ejection process. Furthermore, in the open conformation there is an undefined density which forms a conduit that extends the tube lumen, presumably part of the TMP (**Fig. 4d,e**). Overall, this state reveals the mechanistic basis of a channel formation upon interaction with the lipid bilayer and the initial ejection of the genome.

### The pore forming domains of the TMP

We decided to further study the TMP domains released after contraction and their possible interaction with lipids. AlphaFold2 predicted that the TMP_HBD_ (which is C-terminal to the TMP_TM_, and we previously described interacting with the open tube) can form a globular domain (**Extended Data Fig. 9a**). Based on this prediction, we designed a TMP-truncated construct encoding both the TMP_TM_ and TMP_HBD_ (TMP^S^ (TMP short); TMP (205-518)). Later, we overexpressed and purified the construct from the membrane fraction of the lysate (**Fig. 5a, Extended Data Fig. 9b**). We characterised the sample using single particle cryo-EM, revealing a heterogeneous pool of particles (**Extended Data Fig. 9c,d**). Finally, extensive processing led us to model a 6 Å flexible dimer of TMP^S^ (TMP^S^_2_) (**Fig. 5b, Extended Data Fig. 9e,f**).

**Figure 5.**
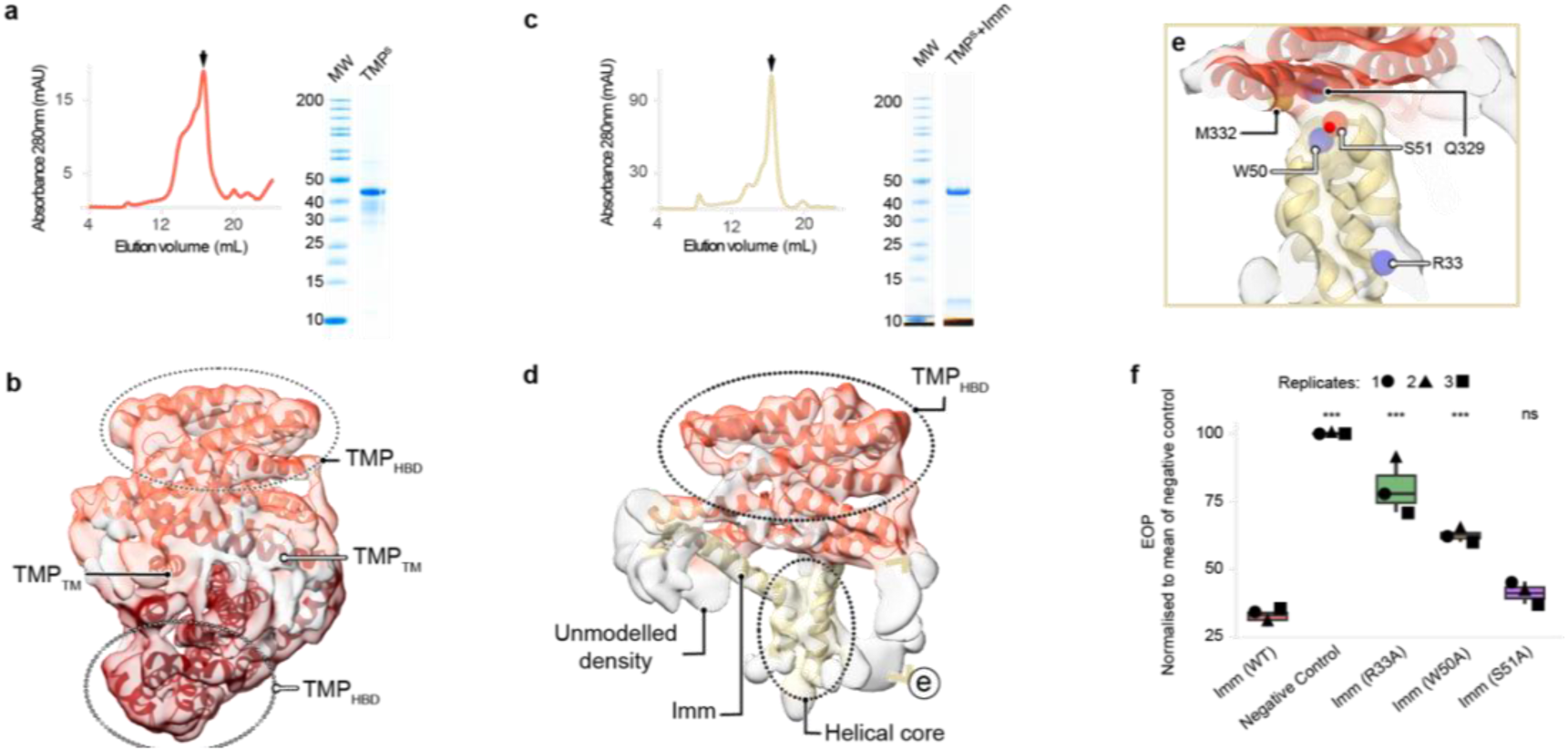
Structure of the TMP^S^_2_ and TMP^S^-Imm complex. **a,** Size-exclusion chromatography profile of purified TMP**^S^** and corresponding cropped SDS-PAGE gel image of the fraction in the arrow. **b,** Cryo-EM reconstruction of the TMP^S^_2_ and the refined model. **c,** Size-exclusion chromatography profile of co-purified TMP**^S^**-Imm and corresponding cropped SDS-PAGE gel image of the fraction in the arrow. **d,** Cryo-EM reconstruction of the TMP^S^_2_-Imm and the refined model. **e,** detailed view of the interface between Imm and the TMP_HBD_, showing the fused density between Imm and the TMP**^S^**. **f**, The effect of Imm mutations on Imm-mediated superinfection exclusion against phage T4 measured in EOP. Data are the mean of three replicates, and each replicate is normalised to the plaque number of the empty vector.

The TMP_HBD_ forms a hydrophilic surface exposed to the solvent, while a hydrophobic patch facing the axis of the dimer harbours the TMP_TM_ (**Extended Data Fig. 9g**). This patch is a conserved region of the protein, where hydrophobic residues are presumably relevant for viral infectivity (**Extended Data Fig. 9h,i**). The presence of a predicted transmembrane helix in the TMP is a common feature among phages from different families, not only for myoviruses but also siphoviruses^6–8^ and bioinformatic prediction of transmembrane segments among a range of phages reinforces this assumption (**Extended Data Fig. 9j**). Overall, these results show the structural domain organisation of a TMP and the amphipathic arrangement of the bundle domain.

Subsequently, to better understand if the TMP is directly involved in genome translocation, we studied if Imm activity requires direct interaction with the TMP. We previously hypothesised that Imm plays a similar role to the immunity proteins of pore forming colicins in blocking the formation of the pore^36,37^. Therefore, we first co-expressed both proteins from the same vector, leading us to co-purify a stable complex of the TMP^S^ bound to Imm from the membrane fraction of the lysate. Afterwards, we performed single particle cryo-EM of the sample to resolve the complex, revealing a more homogeneous set of particles compared to the TMP^S^ alone (**Extended Data Fig. 10a,b**). From this sample, we reconstructed the TMP_HBD_ interacting with Imm embedded in a detergent micelle (**Fig. 5c, Extended Data Fig. 10c**). Based on an AhlphaFold3 prediction, we fitted the helix 1 of Imm, for which the N-terminus is known to be facing the periplasm^30^, and the conserved hydrophobic helical core domain of helices 2 to 4 in the centre of the micelle (**Extended Data Fig. 10d-f**). The loop of Imm connecting helices 2 and 3 faces the hydrophobic patch of the TMP_HBD_ (**Fig. 5d**). The binding of Imm interferes with the interaction between the TMP_HBD_ and the TMP_TM_, which likely attributes for the

TMP_TM_ not being resolved in the cryo-EM density map. Finally, we show that disrupting the interface between the TMP_HBD_ and Imm by mutating Imm (W50A) in the loop decreased the activity of Imm compared to the wildtype (**Fig. 5e,f, Extended Data Fig. 10g,h**). Mutation of Imm (R33A) in the centre of the core helices also negatively impacts the Imm function (**Fig. 5f**). In summary, these data indicate a direct interaction between the TMP and Imm, as well as relevant residues for the superinfection exclusion activity of the protein.

## Discussion

Our cryo-EM analysis of intact phage T4, combined with functional assays, uncovers remarkable details on how this myovirus initiates infection. Building on earlier work on phage T4 structural biology and Imm-mediated exclusion, we reveal insights into the dynamic interplay of the LTFs and the baseplate, and the role of the TMP in the genome delivery. We show that TMP not only defines tail length^5^ but actively regulates DNA translocation: it arrests ejection after contraction and later remodels to form a membrane-spanning conduit. Furthermore, we demonstrate that the superinfection exclusion protein Imm directly binds TMP, blocking this process. Together, these findings provide an important comprehensive structural framework for early infection and membrane penetration by a contractile-tailed phage (**Supplementary Movie 1**).

Host recognition by phage T4 emerges as a dynamic, environmentally responsive process mediated by LTF extension (**Fig. 6a**). Our data identify the KC as a central hub for LTF assembly and retraction. The KC harbours a CBM-like domain that interacts with the whiskers. The charged residues within this domain may confer sensitivity to environmental conditions, such as ion concentration. CBM domains in other phages mediate reversible binding to host LPS^50–52^, suggesting a potential binding role for the KC. We therefore raise the hypothesis that the KC of extended fibres could interact with host LPS, hindering retraction and promoting reversible attachment to the host.

**Figure 6.**
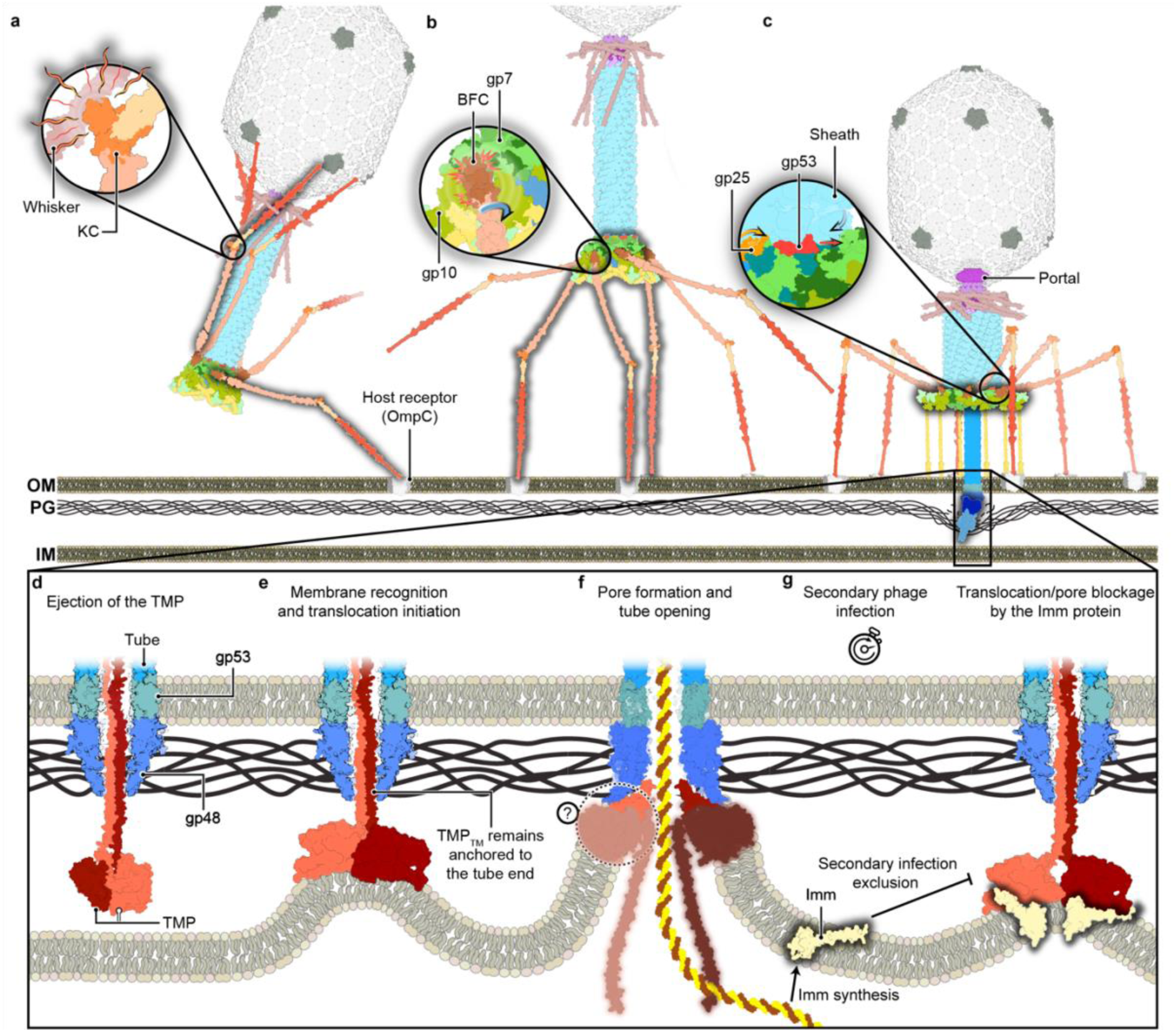
Schematic model of T4 early infection stages and proposed mechanism of genome delivery. **a**, LTF extension and reversible binding to the host. LTF conformational equilibrium can be shifted by environmental sensing of the KC-whisker interaction. **b**, LTF binding triggers baseplate rearrangement. LTF binding to the membrane receptors exerts forces in the baseplate through the BFC to promote peripheral baseplate rearrangement. **c**, Sheath contraction upon irreversible binding of the STFs. Axial expansion of the inner baseplate releases the tube and initiates sheath contraction. The hub punctures the outer membrane (OM) of the host, allowing digestion of the peptidoglycan (PG). **d**, After PG digestion, the bundle domain of the TMP can reach the inner membrane (IM). The TMP conformation is based on the TMP^S^_2_ model. **e**, The TMP recognizes the IM and the TM helices insert into the membrane, initiating translocation of the helices of the bundle domain, bringing the IM closer to the tube. **f**, The TMP and the tube form a conduit for the translocation of the genome into the cytoplasm of the host. **g**, Imm is expressed as an early gene, embedding in the IM and blocking secondary infections. The blurred regions in panels d-g represent the TMP shown only with illustrative purpose and for which structure was not resolved.

Fibre binding appears to exert torque on the baseplate, triggering its rearrangement (**Fig. 6b**). LTFs anchor via the BFC to the docking loop of gp7, which rotates upon extension. Once a number of fibres engage with the host, additional forces such as those exerted during motion of the host may amplify torque on gp7_III_, transmitting the signal to the peripheral baseplate and promoting STF deployment and irreversible binding to the LPS of the host^19,20^. Subsequently, the inner baseplate must disengage from the tube to initiate sheath contraction^18^. In phage T4, we identify the N-terminal loop of gp48 inserted into the baseplate–sheath interface. Its release exposes new binding surfaces on gp25 and gp53, unlocking the sheath layer adjacent to the baseplate and enabling contraction (**Fig. 6c**). Similar interlacing of tube proteins within baseplate architecture occurs in other contractile systems, such as R2 pyocin^28^, likely conferring stability to the pre-contracted state by securing the distal end tube and locking the extended sheath.

The lumen of the tube is filled by the TMP and the genome. Notably, we observe two distinct conformations of the TMP_CL_ interacting with the baseplate hub, which may indicate that the chains are arranged in pairs. Six TMP copies have been reported in mature virions of other phages, including Pam3^53^, Bas63^54^, ur-lambda^10^, and HFTV1^49^, this last one showing a distinct arrangement at the baseplate-hub. Chemical contraction of phage T4 virions revealed partial TMP ejection from the lumen after baseplate-hub detachment, with the ejected part presumably corresponding to the TMP_HBD_ and TMP_CL_ (**Fig. 6d**). About 35% of the initial length of the TMP is retained in the tube lumen, suggesting that the helices arrested by the distal end tube may correspond to the TMP_TM_ region. This would comply with a process where the transmembrane region is only released once the lipid environment is present.

Our data further implicate the TMPs in membrane penetration. Predicted transmembrane segments across diverse phages indicate a conserved membrane-interacting role. Upon release, the TMP recognizes lipid bilayers, consistent with prior observations of lipid recognition by contracted phage T4^23^ and cryo-ET models of tube–membrane interaction^24^. The amphipathic TMP_HBD_ may facilitate membrane curvature toward the tube end^55^ and/or its own translocation (**Fig. 6e**). This process brings the inner membrane closer to the distal end tube, where the TMP_TM_ can be inserted, opening the tube and leading to the translocation of the rest of the TMP and the phage genome (**Fig.6f**).

Finally, we provide compelling evidence for TMP-mediated pore formation by demonstrating its direct interaction with the superinfection exclusion protein Imm. The purified complex reveals a stabilized TMP_HBD_ bound to Imm, suggesting a state in which the interplay between the TMP_HBD_ and the TMP_TM_ to open a translocation conduit for the phage genome is blocked. A similar mechanism has been proposed for gp15 of phage HK97, another superinfection exclusion protein which interacts with the TMP to prevent genome translocation^35^ and therefore indicating a possibly conserved strategy. This resembles the role of the immunity proteins of pore-forming colicins, which inhibit pore formation by blocking the translocation of the pore-forming domain after membrane engagement^36–38^. Colicins also employ helical bundle domains for pore formation^56–58^ and there has been an example showing they can translocate DNA bound to them^59^. These similarities strengthen the case for the TMP acting at the inner membrane and underscore its function as a pore-forming element in genome delivery.

Overall, these findings establish a detailed structural framework for phage T4 infection, from host recognition to genome ejection and its inhibition, and highlight conserved principles of contractile systems. This work provides a foundation for future investigations into phage genome delivery and superinfection exclusion mechanisms.

## Methods

### Bacterial handling and culturing

*E. coli* strains NEB10β, BL21Star(DE3), B (11303), CR63 and K-12 MG1655 were routinely cultured in Lysogeny Broth (LB) under aerobic conditions at 37 °C, unless otherwise stated. LB 1% (w/v) agar plates were used as solid medium and plasmid maintenance was performed using ampicillin at 100 µg/ml, chloramphenicol at 34 µg/ml or kanamycin at 50 µg/ml.

### Plasmid construction

Plasmid cloning involving DNA fragment insertion into the backbone was carried out following the NEBuilder HiFi DNA assembly reaction protocol (New England Biolabs). Site-directed mutagenesis of the plasmids was carried out by PCR amplification with 15 bp of overlapping primers and directly transformed into bacteria or first circularised using the KLD Enzyme Mix (New England Biolabs). *E. coli* NEB10β was used as host for all cloning. All plasmids were confirmed using Sanger sequencing or Nanopore whole plasmid sequencing by Macrogen-Europe.

### Plasmids for CRISPR-Cas engineering

Three different plasmids were designed to target the 5’ region of the *g29*, encoding the TMP, with CRISPR-Cas9 expression using the SpCas9 variant. We designed 20 bp spacers to express the sgRNA: (i) ttcagataataaaccaacac, (ii) cccagataaagtattagaag and (iii) gcagtgagtgatactactgc. We used the backbone of pAH212 for sgRNA expression and the vector DS-SPcas for Cas9 expression^60^.

The homology plasmid for recombination was cloned from the genomic region of bacteriophage T4 DNA. A 658 bp fragment was amplified from the T4 genomic DNA used in this study and cloned into a pUC19 vector. The region amplified was between nucleotides 118,740 to 119,401 based on NC_0008666.4. The resulting plasmid was used as template for the amber mutations on the *g29* codons encoding for residues T77 and T78. The final plasmid was transformed on *E. coli* CR63 amber suppressor strain.

### Plasmids for proteins expression and purification

The vector pET11a was used to clone the sequence encoding for the TMP^S^ bound to a Twin Strep-tag on its C-terminus. This was the vector used for expression and purification of the TMP^S^.

The vector pRSFDuet-1 was used as backbone for *Imm* cloning into the first multiple cloning site and subsequent mutagenesis. Notably, for cryo-EM data collection we purified the 94 residue-long Imm protein starting with an earlier codon than the 83 residues-long reported in the databases (Uniprot P08986). This start codon is present in the reference genome (NC_0008666.4). This was initially done to observe if the extended N-terminal helix could have any structural relevance on the formation of the complex. The reconstruction showed that the extra residues do not participate in the TMP^S^–Imm interface, therefore for the downstream functional assays we used the 83 residue-long protein.

For the co-purification experiments, the sequence encoding the TMP^S^-tag was cloned in the second MCS of the vector pRSFDuet-1 to express both, Imm and the TMP^S^ from the same plasmid.

### Amber *g29* homology plasmids

We designed a plasmid encoding the gp29 carrying silent mutations along its sequence with flanking homology arms. The silent mutations were required to avoid only the repair of the amber mutations and promote whole gene recombination. The gene was synthetised by Twist Bioscience and later we cloned the flanking homology regions (470 and 466 bp upstream and downstream from *g29*, respectively). This vector was used as template for point mutations.

### Bacteriophage propagation and high titre purification

Bacteriophage T4 was obtained from the Félix d’Hérelle Reference Center for bacterial viruses. The phage was purified using PEG precipitation and density gradient method. Phages were propagated in *E. coli* B (11303) (ATCC) at an OD_600_ of 0.3. The cleared lysate was mixed with 10% PEG and 1 M NaCl and incubated overnight at 4°C. The lysate was centrifuged (15,000 *xg*, 1h, 4 °C) and the precipitate was resuspended in minimal amount of SM buffer (100 mM NaCl, 8 mM MgSO_4_, 50 mM Tris-HCl, pH 7.5) for resuspension. The bacteriophage preparation was purified using rate-zonal centrifugation with OptiPrep^TM^ Density Gradient Medium (Sigma Aldrich) density gradient. The OptiPrep^TM^ layers ranging from 50% to 10% in 10% steps were prepared by diluting with SM buffer. Phage sample was applied on the top of the gradient, centrifuged (150,000 *xg*, 18 h, 4 °C) and extracted for dialysis with SM buffer.

### Phage genome sequencing, assembly and annotation

Phage genomic DNA was extracted from culture lysates using the Phage DNA Isolation Kit (Norgen Biotek), according to the manufacturer’s instructions. Library preparation and sequencing were performed by Eurofins Genomics (Illumina NovaSeq platform; 151-bp paired-end reads). Adapter- and quality-trimmed reads provided by the sequencing facility were assembled *de novo* with Unicycler v0.5.1^61^, using the reverse-read (R2) file as input, yielding a single contig circular contig of 168,052 base pairs. Automated genome annotation was performed with Pharokka^62^ (Galaxy v1.3.2+galaxy0) using default parameters. For standardized genome orientation, assemblies were manually linearized in Geneious Prime 2026.0.2 by setting the first nucleotide to the start codon of the predicted terminase small subunit.

The assembled genome was aligned to the reference genome of *Escherichia coli* phage T4 (DSMZ 4505 / Leibniz Institute DSMZ) using MAFFT v7.490^63^. The alignment indicated 100% nucleotide identity across the full length of the genome.

### Chemical contraction of the virions and induction of DNA ejection

Phage contraction was induced by dialysing 100 µl of the purified bacteriophage T4 (4 h, 4 °C) in contraction buffer (3 M urea, 50 mM Tris-HCl pH 7.5, 1 mM MgCl2, 30 μg/ml DNAseI). The contracted phages were diluted 10 fold and pelleted (20,000*g*, 30 min, 4 °C), which kept most of the damaged virions on the supernatant. The pellet was resuspended in SM buffer for vitrification.

Genome ejection was induced by interaction with *E. coli* whole lipids embedded in nanodiscs. The MSP1E3D1 nanodisc purification was performed based on the protocol from Johansen *et al.* (2019)^64^. Nanodisc reconstitution was performed with *E. coli* whole lipids at a ratio of 1:80 (lipids to csMSP1E3D1). 0.11 g of Bio-Beads were added per 200 μl of reconstitution mixture and incubated for 2 h in an orbital shaker at 10 °C. The mixture was separated from the beads by centrifugation and loaded into a size exclusion chromatography column. Nanodiscs were mixed with the pre-contracted virions prior to contraction induction, since this was shown to improve genome ejection rate.

### Mass spectrometry of the virions

100 µL of room temperature 50 mM TRIS pH 8.5 was added to 20 µg of purified T4 phage sample. Following this, 0.5 µg of sequencing-grade trypsin was added and the samples were incubated overnight at 25 °C. Digests were reduced and alkylated by concomitant addition of tris(2-carboxyethyl)phosphine and chloroacetamide to final concentrations of 10 mM, followed by incubation at 30 °C for 30 min. Samples were clarified through 0.45 µm spin filters, and peptides were purified via low-pH C18 StageTip procedure. To this end, C18 StageTips were prepared in-house, by layering four plugs of C18 material (Sigma-Aldrich, Empore SPE Disks, C18, 47 mm) per StageTip. Activation of StageTips was performed with 100 μL 100% methanol, followed by equilibration using 100 μL 80% acetonitrile (ACN) in 0.1% formic acid, and two washes with 100 μL 0.1% formic acid. Samples were acidified to pH <3 by addition of trifluoroacetic acid to a final concentration of 1% (vol/vol), and loaded on the StageTips. Subsequently, StageTips were washed twice using 100 μL 0.1% formic acid, after which peptides were eluted using 80 µL 25% ACN in 0.1% formic acid. The samples were dried to completion using a SpeedVac at 60 °C. Dried peptides were dissolved in 20 μL 0.1% formic acid (FA) and stored at −20 °C until analysis using mass spectrometry.

Samples were analysed on a Vanquish™ Neo UHPLC system (Thermo) coupled to an Orbitrap™ Astral™ mass spectrometer (Thermo). Samples were analysed on 15 cm long analytical columns, with an internal diameter of 75 μm, and packed in-house using ReproSil-Pur 120 C18-AQ 1.9 µm beads (Dr. Maisch). Elution of peptides from the column was achieved using a gradient ranging from buffer A (0.1% formic acid) to buffer B (80% acetonitrile in 0.1% formic acid), at a flow rate of 250 nL/min. Gradient length was 30 min per sample, including ramp-up and wash-out, with an analytical gradient of 20 min ranging in buffer B from 5-38%. The column was heated to 45 °C using a column oven, and ionization was achieved using a NanoSpray Flex™ NG ion source (Thermo). Spray voltage set at 2 kV, ion transfer tube temperature to 275°C, and RF lens to 50%. All full precursor (MS1) scans were acquired using the Orbitrap™ mass analyser, while all tandem fragment (MS2) scans acquired in parallel using the Astral™ mass analyser. Full scan range was set to 300-1,300 m/z, MS1 resolution to 120,000, MS1 AGC target to “250” (2,500,000 charges), and MS1 maximum injection time to 150 ms. Precursors were analysed in data-dependent acquisition (DDA) mode, with charges 2-6 selected for fragmentation using an isolation width of 1.3 m/z and fragmented using higher-energy collision disassociation (HCD) with normalized collision energy of 25. Monoisotopic Precursor Selection (MIPS) was enabled in “Peptide” mode. Repeated sequencing of precursors was minimized by setting expected peak width to 10 s, and dynamic exclusion duration to 7.5 s, with an exclusion mass tolerance of 10 ppm and exclusion of isotopes. MS2 scans were acquired using the Astral mass analyser. MS2 fragment scan range was set to 100-1,500 m/z, MS2 AGC target to “100” (10,000 charges), MS2 intensity threshold to 100,000 charges per second, and MS2 maximum injection time to 5 ms.

### Data analysis

MS RAW data were analysed using MaxQuant software (v.2.5.2.0). Default MaxQuant settings were used, with exceptions specified below. For generation of theoretical spectral libraries, the T4 FASTA database was manually constructed and curated via next-generation sequencing. In-silico digestion of proteins to generate theoretical peptides was performed with trypsin, in semi-specific digestion mode, with minimum peptide length set to 6 and maximum peptide length set to 55. Allowed variable modifications were oxidation of methionine (default), protein N-terminal acetylation (default), deamidation of glutamine and asparagine, and peptide N-terminal conversion of glutamine to pyroglutamate, with maximum variable modifications per peptide set to 3. Second peptide search was disabled. Label-free quantification (LFQ) was enabled, with “Fast LFQ” disabled. Stringent MaxQuant 1% FDR data filtering at the PSM-and protein-levels was applied (default).

A summary of the mass spectrometry results can be found in the Supplementary Table 1.

### Salt dependent fibre conformation imaging

Phage T4 sample at 10^9^ PFU/ml in SM buffer was buffer exchanged to SM buffer supplemented with 350 mM MgCl using 20kDa MWCO cassettes, overnight at 25 °C. The dialysed sample was imaged with TEM as described in the following methods section to analyse LTF extension. The same sample was then dialysed back overnight into SM buffer and imaged with TEM to analyse reversibility of LTF conformation.

### Purification of the TMP construct and Imm protein

*E. coli* BL21Star(DE3) electrocompetent cells were transformed with the pET11a or pRSFDuet-1 vectors harbouring the TMP^S^ or Imm and the TMP^S^ constructs. Cells were grown in Terrific Broth (TB) and induced with Isopropyl β-d-1-thiogalactopyranoside (IPTG) 0.5 mM at an OD_600_ of 1. The temperature was then reduced to 18 °C for overnight induction (18 h). Cells were collected by centrifugation and resuspended in lysis buffer (20 mM Tris-HCl pH 7.2, 300 mM NaCl, 10% glycerol) supplemented with 0.5 mM MgCl2, 1 mg of DNaseI, and EDTA-free protease inhibitor (Thermo Fisher Scientific).

The cells were lysed using an Avestin Emulsiflex C3 homogeniser cooled to 4°C, and un-lysed cells and insoluble debris were removed by an initial centrifugation (8000*g*, 15 min). This method was preferred to osmotic lyses as it helped to remove inclusion bodies. The membranes were then pelleted by centrifugation (30000*g*, 40 min) and resuspended in solubilisation buffer (25 mM Tris-HCl pH 7.2, 150 mM NaCl, 10% glycerol, 1% Lauryl maltose neopentyl glycol (LMNG), 1 mM TCEP, also supplemented with protease inhibitor cocktail). After solubilisation (4 h at 4 °C), the membranes were cleared by centrifugation (100,000 *xg*, 40 min). The supernatant was run over a gravity column with 2 ml Strep-Tactin 4Flow high-capacity resin (IBA), washed with 10 ml Wash Buffer (20 mM Tris-HCl pH 7.2, 300 mM NaCl, 10% glycerol, 0.005% LMNG) and eluted in 12 ml of Elution Buffer (20 mM Tris-HCl pH 7.2, 300 mM NaCl, 10% glycerol, 0.005% LMNG, 10 mM desthiobiotin). Prior to preparation of cryo-EM grids, eluted proteins were concentrated using PES membrane centrifugal filters (Cytiva) and equilibrated into the Final Buffer (20 mM Tris-HCl pH 7.2, 300 mM NaCl, 1 mM TCEP, 0.002% LMNG) over a Superose 6 Increase 10/300 GL size exclusion chromatography column (Cytiva).

### TEM of negative stain samples

For qualitative assessment and analysis of LTF extension, transmission electron microscopy (TEM) of negatively stained samples using 2% uranyl acetate was used. Continuous carbon grids with 300 mesh cooper mesh were glow-discharged (30s, 15 mA, in a Leica ACE 200) before application of 3.5 µl of sample. Afterwards, samples were washed 3 times with the corresponding buffer for each sample, always without detergent, and dried before imaged on a Morgagni 268 268 transmission electron microscope operated at 100 kV (FEI/Philips), equipped with a side-mounted Olympus Veleta camera with a resolution of 2048 × 2048 pixels (2 K × 2 K). Images were recorded using ITEM software.

### Cryo-EM data acquisition

3 µl of the purified phage T4 on different conformations were applied on UltrAuFoil grids (R2/2, mesh 200), previously glow-discharged (30 sec, 10 mA, in a Leica ACE 464 200), and plunged-frozen into liquid ethane, pre-cooled with liquid nitrogen, using a Vitrobot Mark IV (FEI, Thermo Fisher Scientific) with 100% humidity at 4°C. TMP^S^ and Imm-TMP^S^ samples were vitrified on NiTi alloy and UltrAuFoil grids, respectively, mixed with 0.025% CHAPSO prior to vitrification.

Data collection was performed on a Thermo Scientific Titan Krios G2 at 300 kV (Thermo Fisher Scientific). For the pre-contracted and post-contracted phage the microscope was equipped with the Falcon 3EC direct electron detector on linear mode at a pixel size of 1.08 Å/pix, a total dose of 40 and 43 e^-^/Å^2^, respectively, and a defocus range of 0.5 to 2.5 µm. The contracted phage interacting with nanodiscs and the TMP samples were collected on the same microscope but equipped with a Falcon 4i Direct Electron Detector and Selectris X Imaging filter. For the phage it was used a calibrated pixel size of 1.2 Å/pix and a total dose of 40 e^-^/Å^2^, while for the TMP samples it was used a pixel size of 0.728 Å/pix and 50 e^-^/Å^2^. Micrographs were collected using the semi-automated acquisition program EPU (FEI, Thermo Fisher Scientific). Number of micrographs and other values for the different dataset are summarized in Supplementary Table 2.

### Reconstruction of the virion

Micrograph pre-processing was done in a similar way for all the datasets using cryoSPARC v4.3.0 to 4.7.0^65^. Patch motion correction and patch contrast transfer function (CTF) estimation were performed before curation of low-quality data based on ice thickness, CTF values and total motion of the sample.

#### Baseplate

For all the phage samples, manually picked baseplates were used to train a Topaz model^66^, later used for particle picking on the full dataset. Particles were extracted on a 750 px box size and Fourier-cropped to 352 px. Two rounds of 2D classification were followed by an *ab-initio* reconstruction of the baseplate. One round of heterogeneous refinement was used to further curate the particles to separate remaining neck complexes. 3D non-uniform refinement with imposed C6 symmetry was used to achieve high-resolution maps.

3D classification without alignment enclosing the masked baseplate-hub ((gp5-gp27)_3_) was used to separate the particle pool into two classes to break the symmetry mismatch. The particles of the first class were turned 120° around the tail axis to align with those of the second class. The map of the second class was used as input for the 3D non-uniform refinement with imposed C3 symmetry. This allowed to achieve the C3 symmetrised map of the phage baseplate. The peripheral, intermediate, and the inner baseplate were further refined by local reconstruction of these regions. For the intermediate and peripheral baseplate, the local refinement job was run with symmetry expanded particles.

#### Neck and portal

Initial picking of the neck region was performed in the same manner as for the baseplate. A mask was used covering the last two layers of the tail sheath to the portal and 3D non-uniform refinement with imposed C6 symmetry led to high resolution map of the tail-neck-portal complex. The portal showed the lowest resolution, so we performed a local refinement with a mask enclosing the (gp13_12_-gp20_12_) complex to improve the density.

The portal of the pre-contracted virions sample was classified into the retracted and extended crown helices. One third of the portals where in the extended helix conformation, this was attributed to the presence of damages capsids in the pre-contracted sample, which can affect genome packaging pressure inside the capsid. The post-contracted and post-ejected genome states presented only the extended helices.

#### Capsid-portal

To break the symmetry mismatch between the portal-tail (C6) and the capsid (C5) it was first resolved the capsid by imposing C5 symmetry and later 3D classification without alignment of the masked portal-neck complex into 6 classes. Particles from the first 5 classes were rotated around the tail axis to match the position of the 6^th^ class. Refinement of the particles still showed a poorly defined density for the tail–portal region, so we further classified the neck–portal. This led to further classification of the particles into two classes, which led to the final maps of the portal–capsid without symmetry-imposed maps in the two different assembly conformation, where the tail–portal was displace 4° around the tail symmetry axis between both classes.

The pre-contracted phage resulted in the best map for the building of the asymmetric adaptor loops, therefore is included in this study. This reconstruction was further used for atomic modelling. The post-contracted phages mixed with nanodiscs had ejected most of their genomes, so the capsids were used to resolve the Ip3 binding to the MCP. The mask used for refinement of the Ip3 also covered the adjacent MCPs, as a mask only including the Ip3 density did not provide optimal results.

#### Long tail fibres

Fibres were resolved from symmetry expanded pre-contracted particles. The proximal rod was resolved by shifting the box centre from the baseplate, while the distal rod was reconstructed through shifting from the neck. Fibre occupancy was assessed using 3D classification without alignment, followed by local refinement to address flexibility in the proximal region. The proximal rod was divided into P1, P2, and P3-5 for local refinements. The KC was local refined enclosing part of the proximal and distal rods.

The pool of wedges missing the retracted fibres was further studied to determine BFC orientation. A mask was used to cover the extended BFC and perform 3D classification over this. This led to the classification of three states, one fully extended conformation and two at intermediate angles of the BFC.

#### Distal end tube (gp48)_6_

The distal end tube on the post-contracted particles was resolved by iteratively shifting the baseplate coordinates towards the end of the tube. Baseplates of contracted particles were symmetry expanded six times around the symmetry axis and coordinates were displaced towards the (gp48)_6_ end of the tube. Following each coordinate shift, particles were re-extracted and locally refined to correct for tube bending by limiting the rotation and shift search (15° and 9 Å respectively) and gaussian prior (5° and 3 Å) to avoid duplication of particles. The final tube tip was extracted with a box size of 480 or 352 px for the phage only and phage mixed with nanodiscs, respectively. Extensive 3D classification without alignment was performed to curate and resolve heterogeneity of the particles. This led to the reconstruction of the distal end tube interacting with the TMP in the post-contracted virions and the three different distal end tube conformations on the post-ejected genomes virions.

#### DNA densities

The DNA-refined region was obtained by classification of the tube lumen density of symmetry-expanded pre-contracted particles at the tube-neck interface. Symmetry expansion six times around the symmetry axis was required to achieve enough signal for classification. A tight mask was applied for classification of the lumen density. The best class was local refined to study the DNA-protein interface.

Post-contracted particles before genome ejection were used to resolve the DNA-protein interface at the baseplate level. The DNA refined region was performed by classification of the tube lumen density of symmetry expanded particles of the baseplate. This was done as described previously for the lumen density near the neck of the pre-contracted virions.

### Reconstruction of TMP^S^ and TMP^S^-Imm complex

Micrograph pre-processing was done in a similar way for all the datasets using cryoSPARC 4.6.2. Patch motion correction and patch CTF estimation were performed before curation of low-quality data based on ice thickness, CTF values and total motion of the sample.

The particles were picked using blob picker job followed by 2D classification to discard ice contamination, evident noise classes and other junk. 2D classes resembling possible particles were used for ab-initio models setting the highest resolution to 9 Å. Iterative rounds of heterogeneous refinement were used to clean up the dataset and classify the different conformations. Afterwards, non-uniform refinement was directly used, followed by 3D classification without alignment to select the best particles.

For the TMP^S^_2_, particles were exported to Relion 5 to perform 3D refinement using Blush^67^. 3D refinement, post-processing and CTF refinements were used to improve the quality of the map.

### Atomic model building using cryo-EM maps

For the baseplate structures, the published models PDB 5IV5 and 5IV7^18^ were used as starting models for the pre-contracted and post-contracted states, respectively. Remaining proteins were manually built. The intermediate baseplate reconstruction of the hexagonal baseplate was used as reference for the refinement of that region of the inner and peripheral baseplate. Each local model was used as reference for the whole baseplate. For the refinement of the extended BFC, the hexagonal baseplate was fitted into the density for the extended BFC and trimer was fitted into the density using Isolde and applying restraints. The docking loop of gp7 was moved together with the BFC, as it is expected from the star-shape state. The domain P1 of the LTF from the pre-contracted virion, retracted state of the fibres, was used as initial template for the LTF in the peripheral wedge of the star-shape baseplate.

In the pre-contracted virion, the TMP chains built based on the initial models based on the sequence fitted by ModelAngelo^68^.

For figure representation, gp5.4 was rigid body fitted into one of the three possible position from the model PDB 4KU0^69^.

For the portal and capsid, the model 6UZC^70^ was initially used. The local refinement of the portal was used to refine the portal and neck adaptor proteins. This model was later used as reference for the complete neck–portal model refinement.

The distal end tube was first modelled in the pre-contracted state reconstruction. Later, the refined model was used as initial template for the modelling of the distal end tube conformations of the post-ejected genome virions, since the resolution allowed confident refinement of the model. Finally, conformation 3 of the distal end tube was fitted and used as initial model for the building of this complex in the post-contracted virion state. The central density of the TMP hexamer was not built since the reconstruction did not allow confident tracing of the main chain for the α-helices. Isolde was used to fit the α3 and the turn-helix-turn fold, relevant segments of this model that were refined into the density without ambiguity.

Ip3, TMP, Imm, gp35 and gpwac were predicted using AlphaFold2 and AlphaFoldMultimer^71^ (**Extended Data Fig. 11a**) and fitted as rigid bodies in the cryo-EM maps using ChimeraX^72^. AlphaFold3^73^ was used for the prediction of the (TMP-gp48)_6_. RBPseg^74^ was used for the prediction of the (gp37)_3_. The high-resolution model of the LTF (gp34)_3_ tower domain of model PDB 5NXH^75^ was used to apply constrains to the LTF_P3-5_ model refinement. The refinement of the low resolution models of the TMP^S^_2_, TMP-Imm, (gp34)_3_-gp35-(gp36)_3_, Ip3 and distal rod of the LTF was performed in Isolde^76^ using the prediction for reference restraints.

The prediction for the TMP^S^-Imm complex using AlphaFold3 did not resemble the experimental density, therefore each component was predicted separately, which allowed confident fitting of Imm into the density. ISOLDE^76^ was initially used for refinement of the predictions into the densities (**Extended Data Fig. 11b**). Iteratively, real space refinement in Phenix^77^ and manual correction using ISOLDE or Coot^78^ were used to improve the models. Phenix was finally used to assess the geometry of the models and to obtain the map to model FSC curves.

For all datasets, the number of videos, the number of particles used for the final refinement, map resolution and other values during data processing are summarized in the Supplementary Table 1.

### Bacteriophage mutagenesis

Bacteriophage T4 genomic mutant for the gp29 amber mutation^79^ was carried out using homologous recombination followed by Cas9 selection. Three different PAM sites were chosen on the 5’ coding site of *g29*, from which one of them showed a 10^3^-fold reduction of plaques: spacer sequence 5’-gcagtgagtgatactactgc. *E. coli* CR63 was transformed with the plasmids harbouring the DS-SPcas^60^, the sgRNA, and the homology template, harbouring the *g29* with amber mutations on T77 and T78. We performed a one-step assay of genome editing, where recombination and selection occur in the same culture. The infected culture was centrifuged (10,000*g*, 2 min) and the pellet resuspended in 100 µl of M9 media. The sample was serially diluted and each dilution used for a plaque assay using CR63 without plasmids for plating. Single plaques were chosen and plated again on CR63 carrying the DS-SPcas and sgRNA expression plasmids.

From the second round of selection, only two plaques were isolated that could grow on CR63 but not on MG1655. The homology region including the mutations of each phage was PCR amplified and Sanger sequencing to confirm the sequence of the amber mutation.

### Recombination assays of *g29*

*E. coli* K-12 MG1655 chemically competent cells were transformed with each one of the plasmids harbouring the gp29 mutants, encoding the *g29* with silent mutations and codon substitution for each mutant. Overnight cultures were 100 times diluted and grown in presence of antibiotic to OD_600_ of 0.3. Bacteria were mixed with top agar layer (LB 0.5% agar) and amber mutant phage T4 was spread on it. From this step, single plaques were spread into MG1655 without any plasmid to isolate individual mutants that were confirmed by PCR, amplifying their genomic region including the flanking homology arms and Sanger sequencing was used to confirm the point mutants.

Recombination efficiency was studied by plating sequential dilutions of the amber mutant phage T4 into the top agar of MG1655 harbouring each plasmid. Recombination efficiency was addressed as reduction on efficiency of plating (EOP).

The statistical analysis was performed as follows. Three technical replicates were averaged within each biological replicate (n = 3 biological replicates per mutation). Plaque counts were log10-transformed (log10[x + 1] for zero counts). Data were analysed by one-way ANOVA, and Dunnett’s test^80^ was used for multiple comparisons to the positive control. Statistical significance was defined as adjusted p < 0.05. Analyses were performed in R (v2025.05.1+513) using the multcomp package for Dunnett’s test. The plaque count can be found in the Supplementary Table 3.

### Imm functional plaque assay

Overnight cultures of BL21Star(DE3) harbouring each one of Imm-encoding plasmids were diluted 100 times on LB with kanamycin. Induction was carried out immediately after dilution at 30 °C for 3 h with 0.05 mM IPTG. Higher concentrations of IPTG showed impaired cell growth, probably due to toxicity^81^. After 3 h, appropriate phage T4 samples were mixed with 200 µl of each bacterial culture. The mixture was incubated for 10 min at 25 °C. Each sample was mixed with the top agar and let dry. The plates were incubated overnight at 30 °C and plaques were manually count.

Statistical analysis was done as follows. Technical replicates were averaged within each biological replicate (n = 3 biological replicates per cell type). Plaque counts were normalized within each biological replicate by scaling to the mean of the negative control set to 100. Data were analysed by one-way ANOVA, and Dunnett’s test^80^ was used for multiple comparisons to the positive control. Statistical significance was defined as adjusted p < 0.05. Analyses were performed in R (v2025.05.1+513) using the multcomp package for Dunnett’s test. The plaque count can be found in the Supplementary Table 4.

### Bioinformatic analysis

#### Sequence alignment and analysis

Protein sequences were analysed for homology using the BLASTP^82^. The non-redundant (nr) protein database from the NCBI was employed as the search dataset^83^. Each query protein sequence was used as input against the database using the BLOSUM62 scoring matrix. Results with percentage identity >30% and E-value <1e-5 were used for Imm to assert conservation. Should be noted that despite the high E-value all the hits were annotated as superinfection immunity proteins. For the TMP, identity percentage >30% and E-value <1e-10 were uses instead.

Filtered hits were aligned using ClustalOmega via the MPI Bioinformatic Toolkit^84^. These alignments were used as input on ConSurf^85^ to plot the conservation profile on the atomic models.

#### Transmembrane sequence prediction on tape measure proteins

Pre-annotated TMP sequences from the BASEL collection^86^, together with additional selected phages, were retrieved from UniProt. TMP orthogroups were identified using PHROGS^87^, and the corresponding pre-computed multiple sequence alignments (MSAs) were employed to construct hidden Markov model (HMM) profiles with HMMER v3.3.2 (hmmbuild)^88^. A comprehensive HMM database comprising all TMP PHROGS orthogroups was generated with hmmpress, and these profiles were subsequently used to classify the selected TMP sequences into orthogroups with hmmsearch.

Prediction of transmembrane regions was performed with DeepTMHMM v1.0^89^, TMBed v1.0.2^90^, and InterPro v106.0^91^ (incorporating TMHMM and Phobius). Phylogenetic reconstruction of the selected TMP sequences was carried out with IQ-TREE v3.0.1^92^. Multiple sequence alignment was generated with Clustal Omega^93^ v1.2.4 and trimmed with trimAl v1.5^94^; the resulting alignment served as input for IQ-TREE (iqtree -s input.aln -nstop 500 -bb 1000 -m LG+G4 -nt 4). The final phylogenetic tree was visualized using pycirclize v1.10^95^.

#### Structural analysis

We used Foldseek Search Server^96^ to search for similar structures of gp13 and two domains of the kneecap, ILEI/PANDER-like and CBM-like.

Protein structure comparison of the CBM-like domain of the KC and the CBM161-1 [PDB: 3OEB] and CBM22-2 [PDB: 4XUT] was done using TM-align^97^.

For structure comparison between the pre- and post-contracted baseplate conformations the gp53 copies were used for alignment.

### Figure Preparation

Structures were visualised and figures prepared using ChimeraX^72^, Adobe Illustrator, and Adobe Photoshop. Videos were prepared using Adobe Premiere. For the Extended Data Movie S1, the capsid of the phage was derived from the PDB 6UZC^98^.

## Data availability

The cryo-EM maps and the corresponding atomic coordinates have been deposited in the Electron Microscopy Data Bank (EMDB) and in the Protein Data Bank (PDB), respectively. The accession codes are listed as follows. Pre-contracted state: tail EMD 56476, capsid EMD 56658, baseplate EMD 56876 PDB 28VE, inner baseplate EMD 56838 PDB 28UO, intermediate baseplate EMD 56770 PDB 28RJ, peripheral baseplate EMD 56767 PDB 28RG, neck and portal EMD 56768 PDB 28RH, DNA-terminus EMD 56660, portal (withdrawn crown) EMD 56762 PDB 28RA, portal (extended crown) EMD 56774 PDB 28SFretracted LTF EMD 56661, proximal rod of the LTF EMD 56757 PDB 28QV, domain P1 of the LTF EMD 56758 PDB 28QW, domain P2 of the LTF EMD 56759 PDB 28QX, extended state of the BFC EMD 56892 PDB 28VO, intermediate position 1 of the BFC EMD 56670, intermediate position 2 of the BFC EMD 56671, domain P3 to P5 of the LTF EMD 56760 PDB 28QY, kneecap of the LTF EMD 56875 PDB 28VD, distal rod of the LTF EMD 57037, neck and capsid assembly 1 EMD 56662, neck and capsid assembly 2 EMD 56896 PDB 28VX; Post-contracted state: tail EMD 56672, capsid EMD 56673, neck and portal EMD 56877 PDB 28VF, DNA terminus EMD 56674, portal (extended crown) EMD 56865 PDB 28UZ, distal end tube EMD 56744 PDB 28QF; Post-ejected genome: tail EMD 56675, capsid EMD 56676, inner baseplate EMD 56868 PDB 28VC, peripheral baseplate EMD 56756 PDB 28QU, domain P1 of the LTF EMD 56754 PDB 28QR, STF EMD 56743 PDB 28QD, neck and portal EMD 56704 PDB 28PD, portal (extended crown) EMD 56694 PDB 28OX, distal end tube in open conformation EMD 56708 PDB 28PH, distal end tube alternative conformation 2 EMD 56705 PDB 28PF, distal end tube alternative conformation 3 EMD 56719 PDB 28PO, internal protein III EMD 56680 PDB 28ON; recombinantly expressed: tape measure protein dimer EMD 56678 PDB 28OK, tape measure protein in complex with immunity protein EMD 56679 PDB 28OL. The raw cryo-EM datasets used in this study have been deposited in the Electron Microscopy Public Archive (EMPIAR): (ongoing process). The mass spectrometry proteomics data have been deposited to the ProteomeXchange Consortium via the PRIDE partner repository with the dataset identifier PXD072569.

## Supporting information

Summary of phage T4 early infection

## Acknowledgements

We thank Ján Bíňovský for his support in structural model refinement. We thank Blanca Lopez Mendez from the Protein Production and Characterization platform for her support in mass spectrometry.

The Novo Nordisk Foundation Center for Protein Research is supported financially by the Novo Nordisk Foundation (NNF14CC0001 and). N.M.I.T. acknowledges support from an NNF Hallas-Møller Emerging Investigator grant (NNF17OC0031006), an NNF Hallas-Møller Ascending Investigator grant (NNF23OC0081528), an NNF Project grant (NNF21OC0071948) and an LF Ascending Investigator grant (R434-2023-289). N.M.I.T. is also a member of the Integrative Structural Biology Cluster (ISBUC) at the University of Copenhagen.

## Author contributions

A.R.-E. and N.M.I.T conceived the project. A.R.-E. performed phage T4 purification and cryo-EM sample preparation. A.R.-E. collected cryo-EM data with assistance of T.P. and N.S. I.A.H. performed mass spectrometry and analysed the data, in consultation with M.L.N. A.R.-E. processed the cryo-EM data of phage T4 and determined the structures of this study, with assistance of L.M.-A. L.M.-A. performed the fibre extension experiments, with assistance of A.R.-E. A.R.-E. and L.M.-A. performed the cloning of the plasmids in this study. A. R.-E. and N.R.R. expressed and purified the TMP^S^ and TMP^S^-Imm constructs, with assistance of M.S. and H.H. A.R.-E. purified the phage genome and D.P. performed the genome assembly of the phage used in this study. A.R.-E. performed the mutagenesis of phage T4 amber mutant and the assays of the recombinant phages and the activity of Imm, with assistance of A.H. V.K.-S. performed the TMP transmembrane sequence analysis with assistance of A.R.-E. V.K.-S. and A.R.-E. performed the bioinformatic analysis. N.M.I.T. acquired the financial support for the project. A.R.-E. wrote the manuscript, and prepared the figures and movies, with input from all authors. All authors contributed to the revision of the manuscript.

---

**Extended Data Figure 1.**
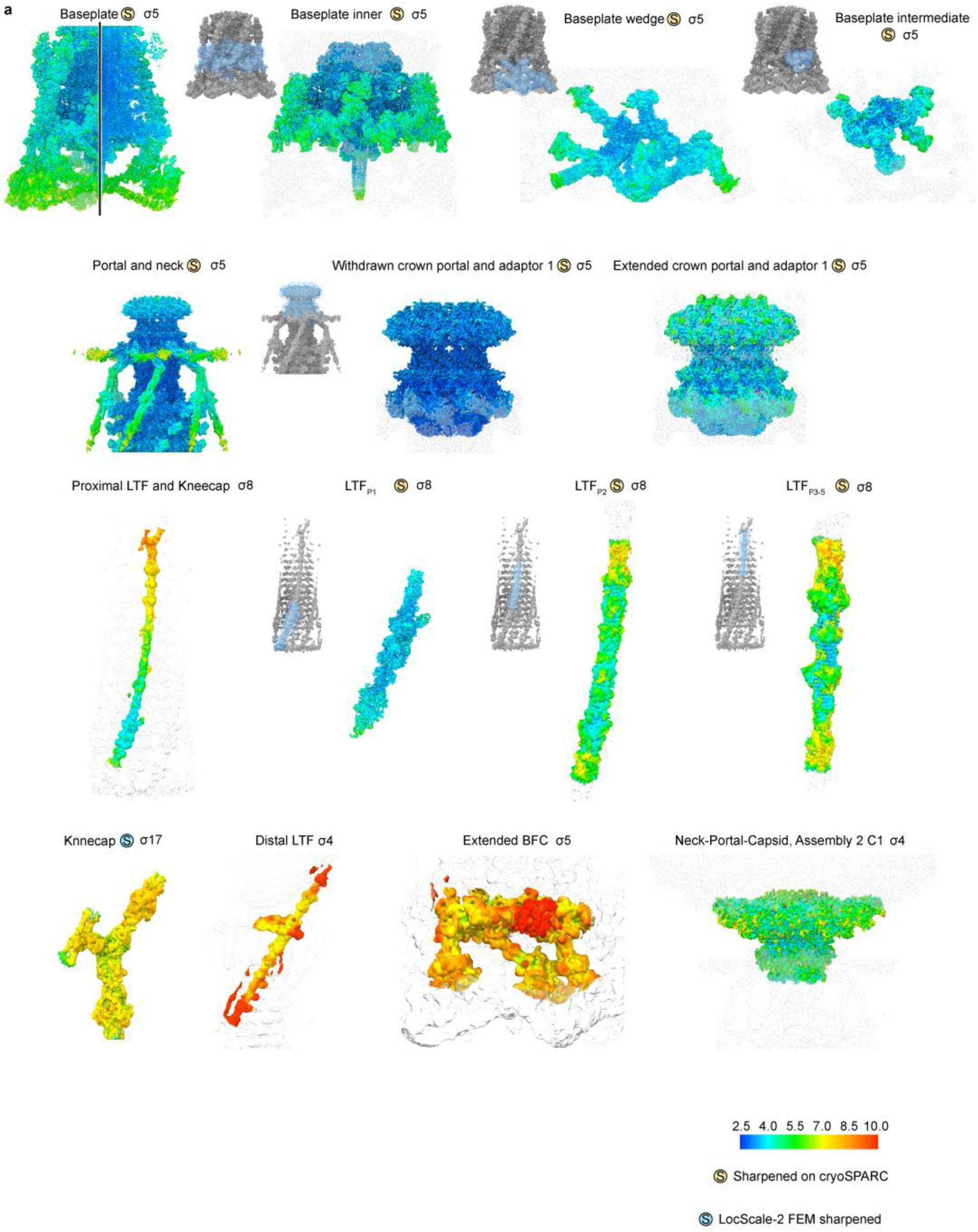
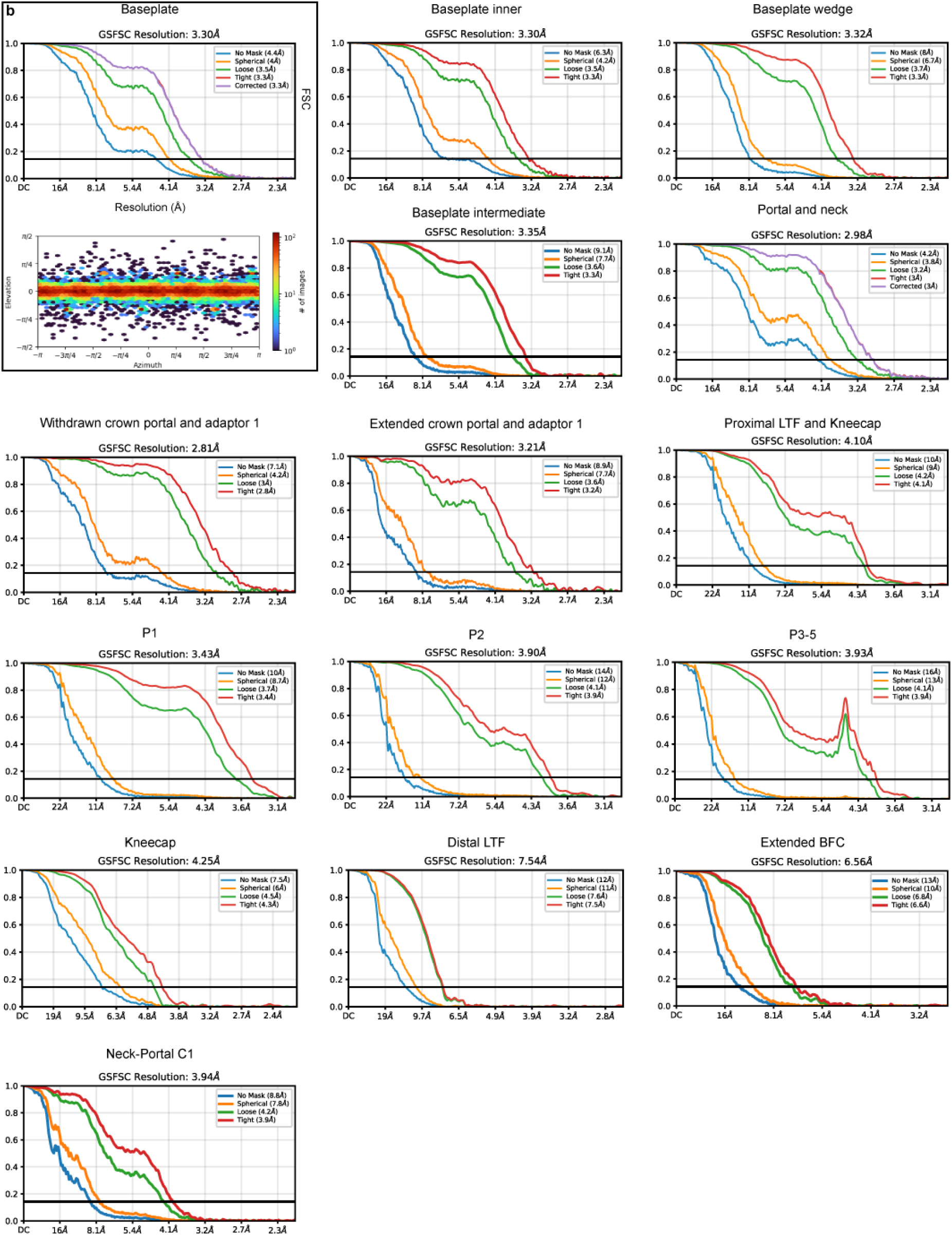
Cryo-EM reconstruction summary of pre-contracted virion. a, Local resolution estimation of the reconstructions used for model building of the pre-contracted virion. **b**, FSC curves of the reconstructions of the pre-contracted virion.

**Extended Data Figure 2.**
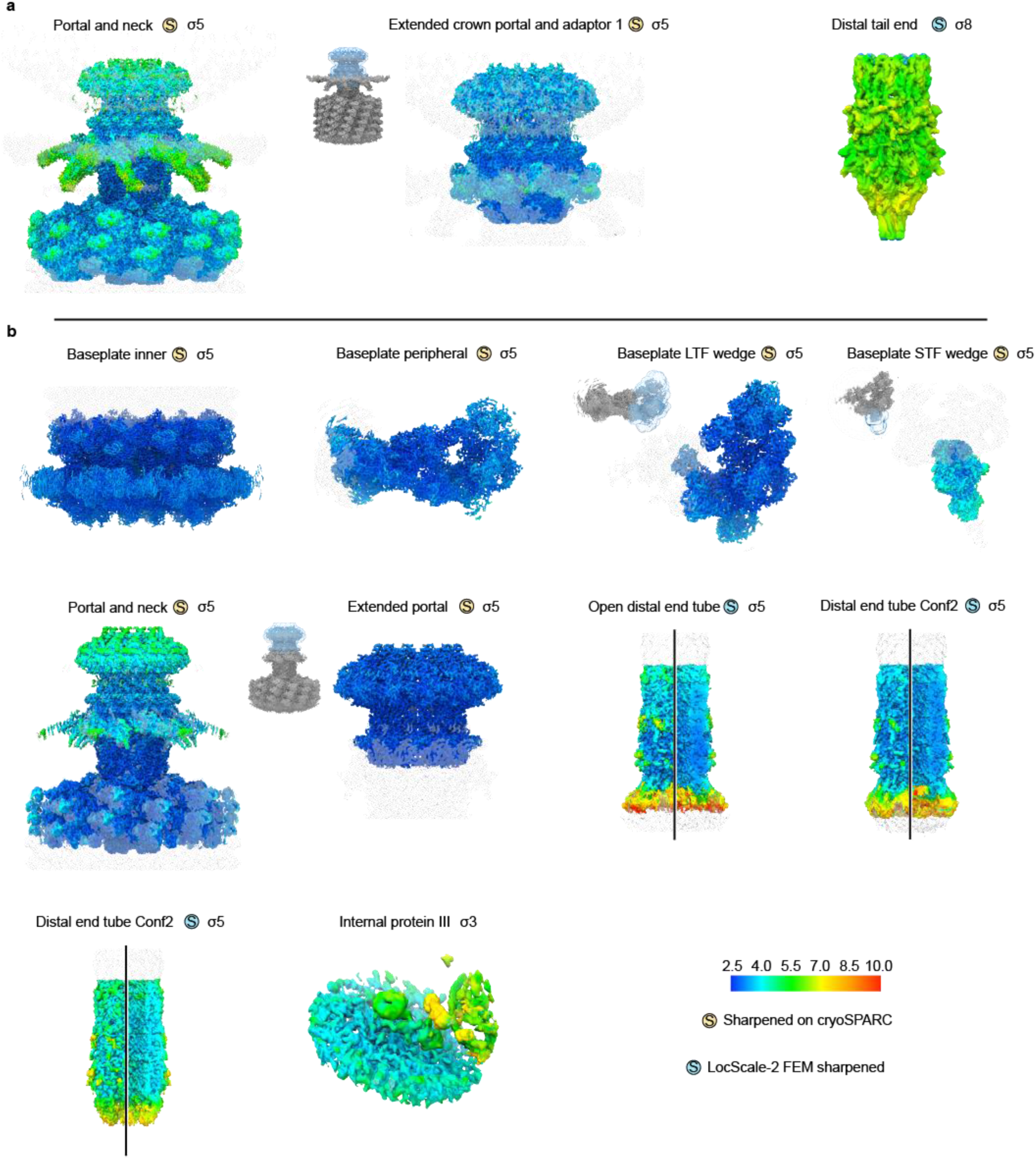
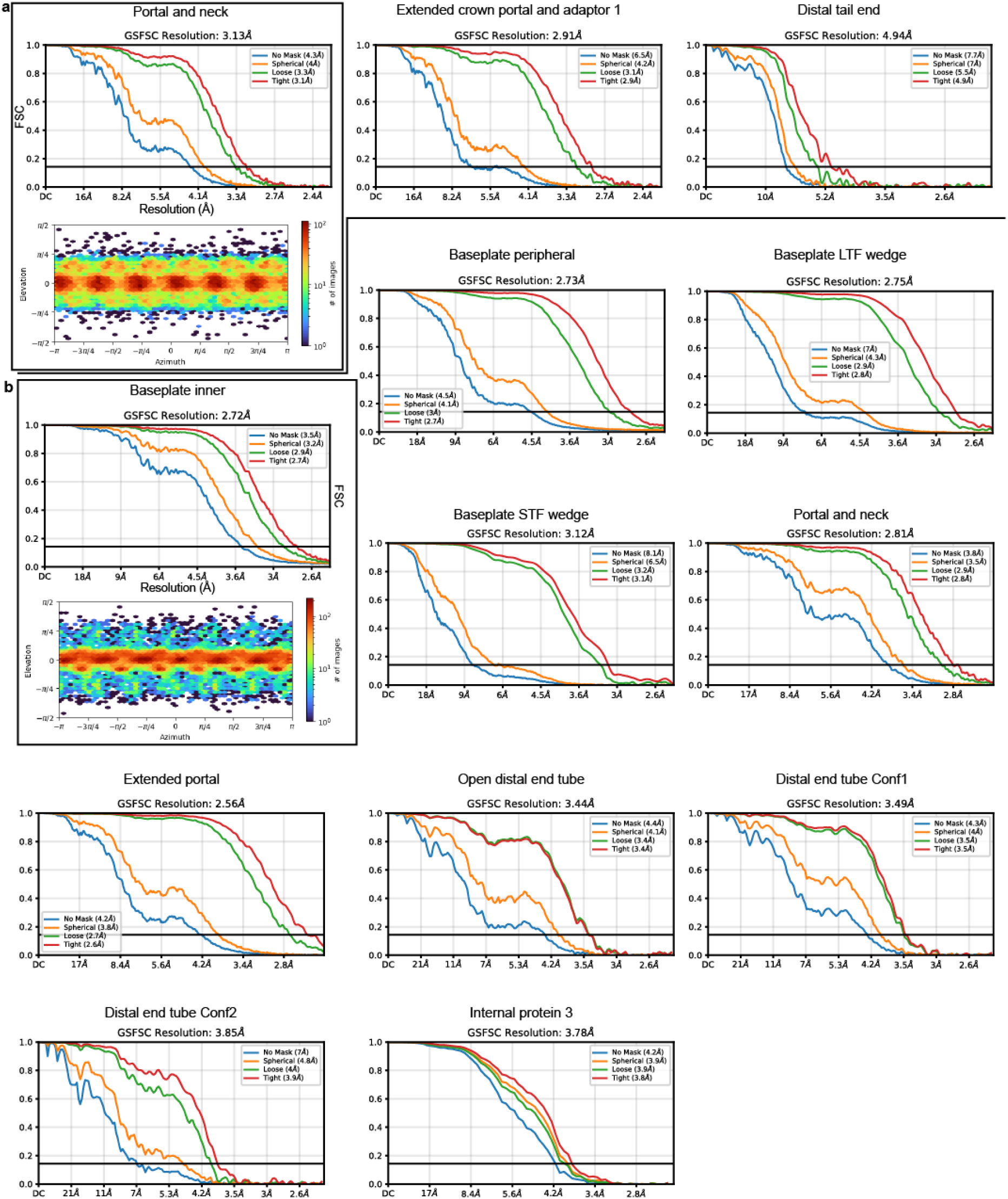
Cryo-EM reconstruction summary of post-contracted virions. and post-ejected genome virion and FSC curves of the reconstructions of the post-contracted virion and post-ejected genome virion. a, Local resolution estimation of the reconstructions used for model building of the post-contracted virion. **b,** Local resolution estimation of the reconstructions used for model building of the post-ejected genome virion. **c,** FSC curves of the post-contracted virion reconstructions. **d,** FSC curves of the post-ejected genome virion reconstructions.

**Extended Data Figure 3.**
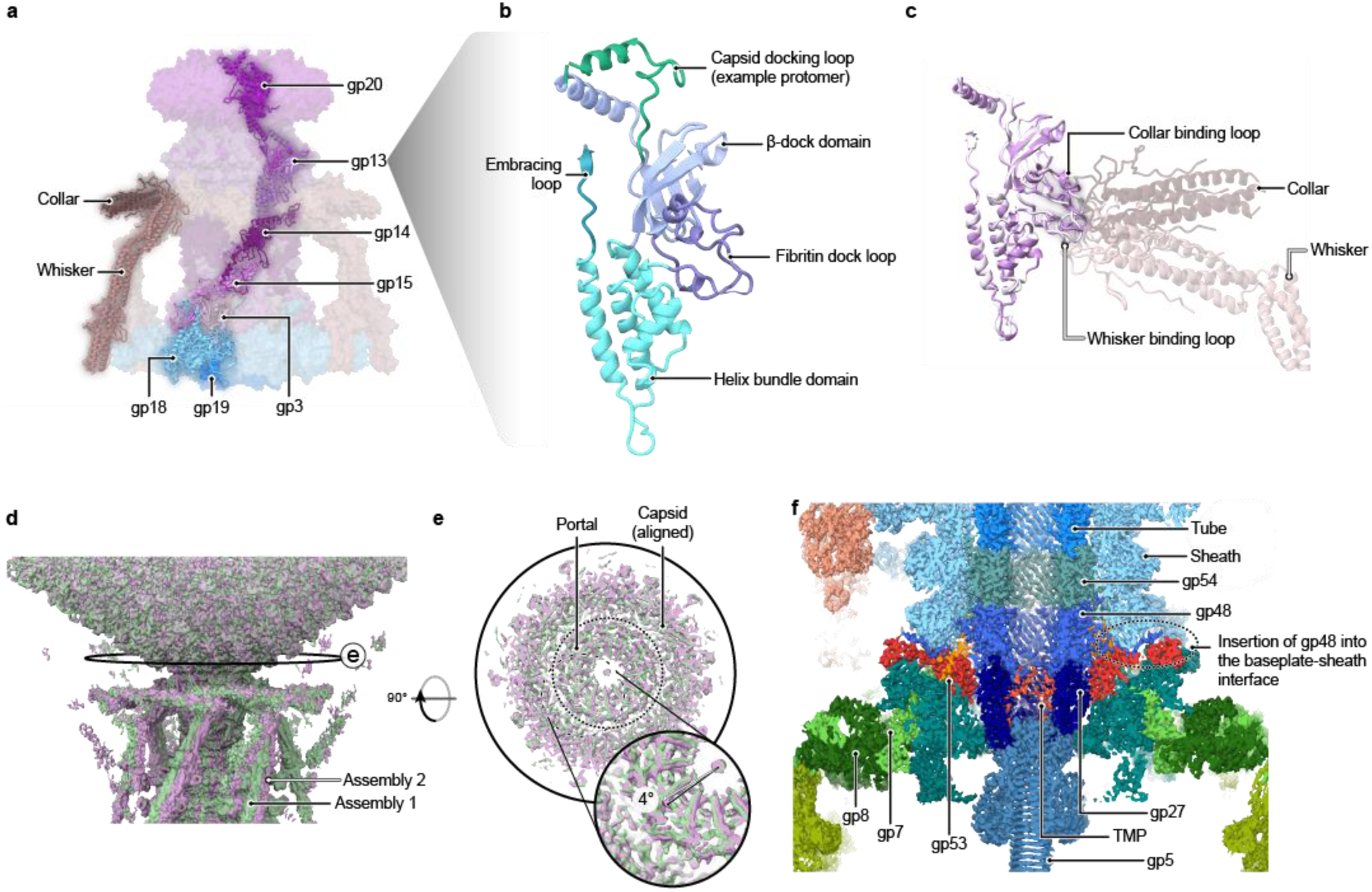
Phage T4 virion architecture. a, Organisation of the neck–portal complex of phage T4. **b,** Structure of a representative monomer of gp13, the neck adaptor. The domains are subdivided based on the gp11 of phage SU10^40^ and gp81 of phage GP4^41^, which were matched by FoldSeek with E-values of 6.11e-2 and 1.03e-1 respectively. **c,** Comparison of one gp13 monomer binding the collar and another binding the whisker. **d,** Alignment of the cryo-EM reconstruction of the two tail-capsid assemblies by alignment of the capsid density. **e,** Clipped top-view of the same alignment as in panel d. **f,** Sliced cryo-EM reconstruction of the inner and intermediate baseplate, and distal tail end.

**Extended Data Figure 4.**
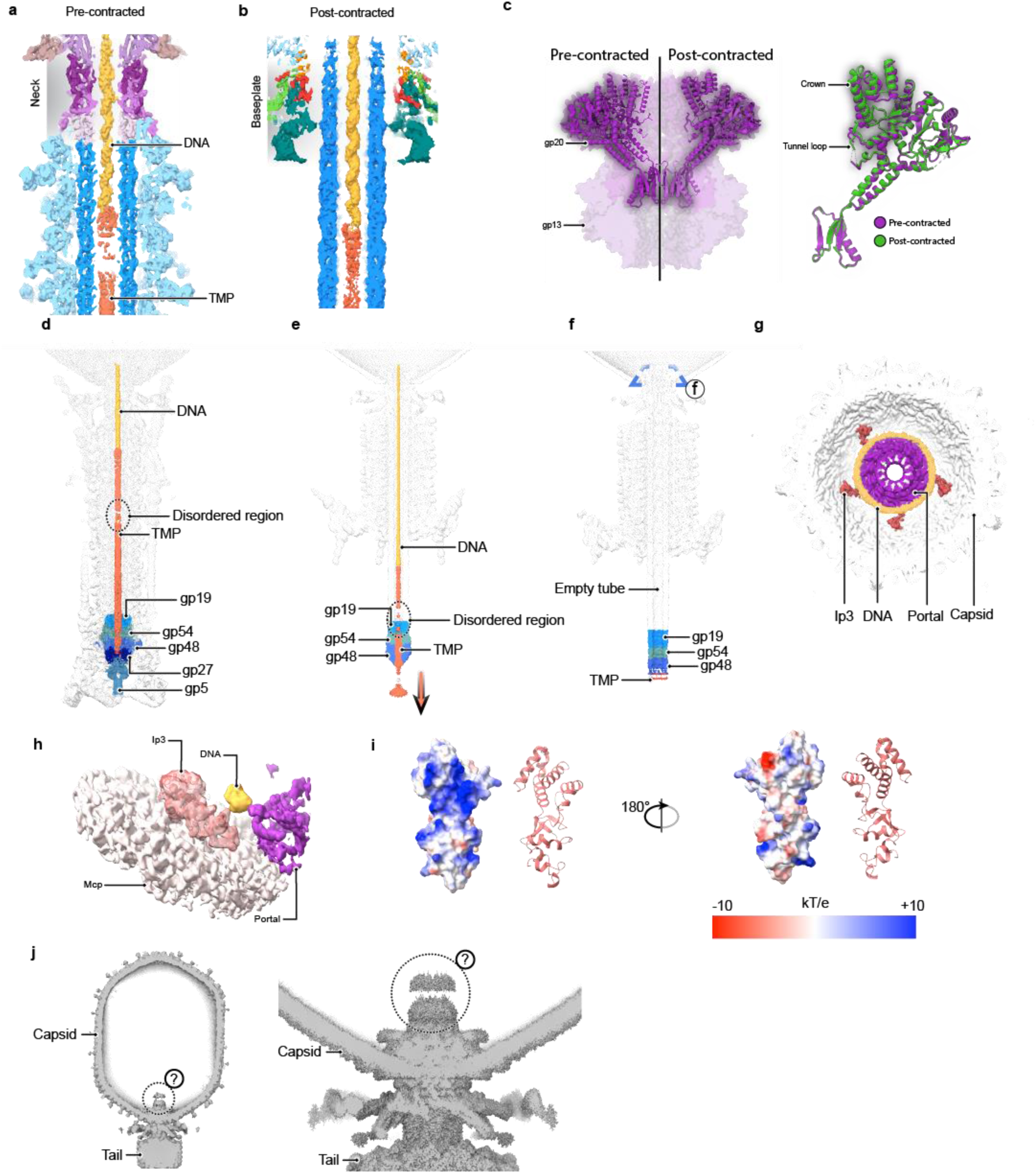
**Overview of genome delivery process**. **a,** Sliced cryo-EM map of the pre-contracted virion at the tube proximal end, where the genome terminus binds to the TMP. **b,** Sliced cryo-EM map of the post-contracted virion at the level of the baseplate, where the genome terminus is retained after contraction. **c,** Comparison between the pre- and post-contracted conformations of the portal protein gp20 and alignment of both portal states. **d**, Sliced cryo-EM map of complete tail and neck of phage T4 in pre-contracted state, where DNA, TMP, baseplate proximal tube, and spike are coloured. **e**, Post-contracted virion state represented as in panel a. Arrow indicates ejection of the TMP. **f**, Post-ejected genome virion state in presence of nanodiscs, represented as in panel e. **g**, View of the portal in the post-ejected genome capsid, showing the density of DNA around the portal and the Ip3 copies (Ip3, internal protein III). **h,** Cryo-EM reconstruction of the local refinement of the Ip3, refined AlphaFold3 prediction fitted in the density. **i,** Surface representation of the Ip3 coloured based on electrostatic potential. **j,** Reconstruction of the capsid without symmetry imposed (*left*) and neck-portal with six-fold symmetry imposed (*right*) at low threshold to show the unresolved density above the portal.

**Extended Data Figure 5.**
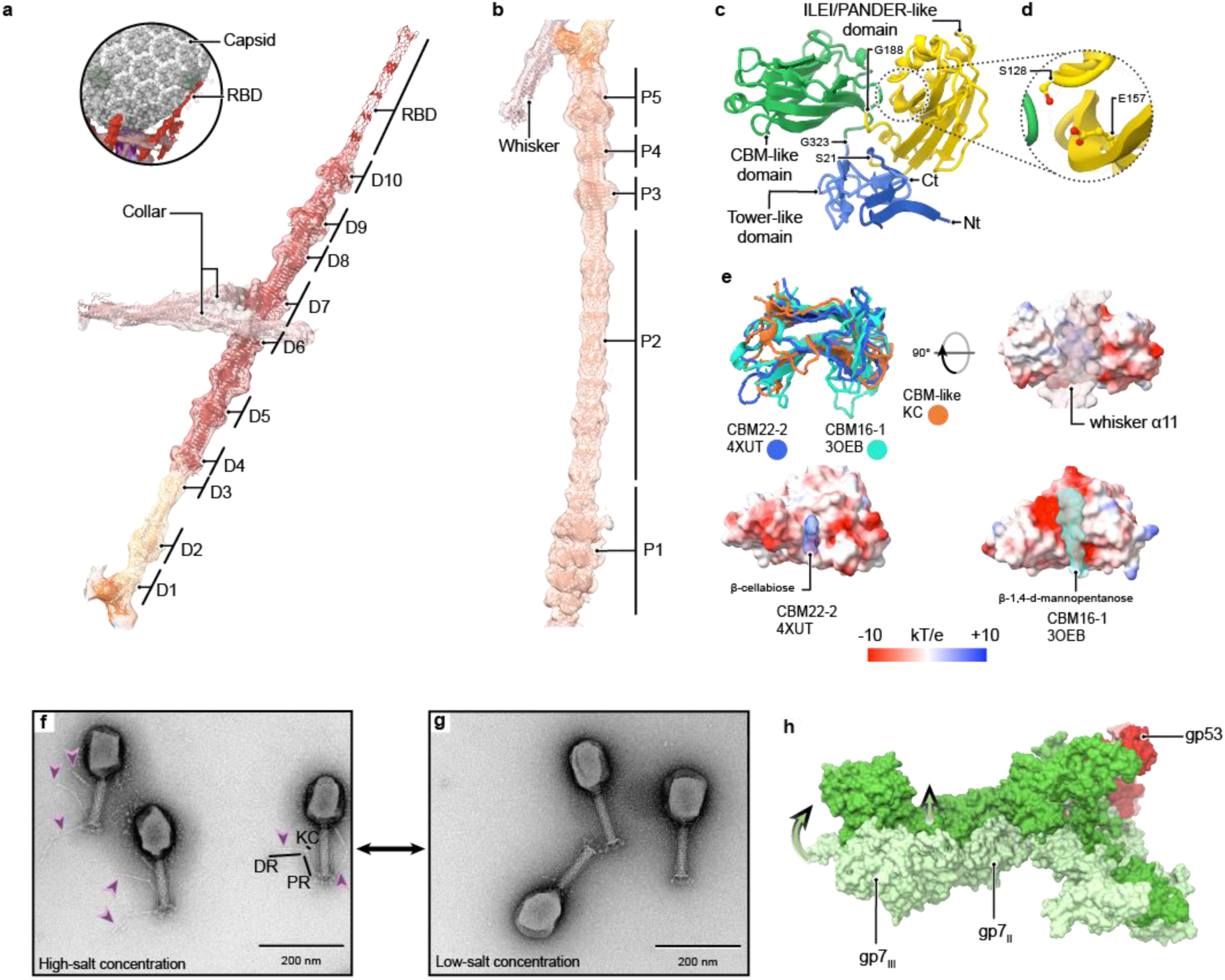
Fibre architecture and extension. **a**, Distal rod, kneecap and collar fibritins cryo-EM map and the refined model fitted. Two collar fibritins clamp (gp37)_3_ in between domains 6 and 7. The RBD could not be resolved for all the distal rods as result of the different bends due to the capsid, but one RBD was bound to the capsid and present on the capsid refinement (*top left*) (RBD, receptor binding domain) **b**, Proximal rod, kneecap and whisker fibritin cryo-EM map and the refined model fitted. **c**, Domain distribution of the kneecap protein. **d**, Detailed view of the conserved hydrogen bound in the ILEI/PANDER-like domain^46^. **e**, Aligned structure of the CBM-like domain of the kneecap to the CBM22-2 and CBM16-1. Each structure is also surface represented facing the groove, where the ligand is found (the whisker α11 for the kneecap) and coloured based on electrostatic potential. The TM-score for the CBM22-2 is 0.712 and RMSD=2.64 Å for 115 residues, and for CBM16-1 is 0.729 and RMSD=2.16 Å for 109 residues. **f**, TEM image of negative stained phage T4 sample in high-salt buffer. Purple arrows indicate extended LTF. **g**, Same sample as imaged in panel g, but after buffer exchange to low-salt buffer. **h**, Gp7 conformational change from the pre-contracted baseplate and the post-contracted state (light and darker colours respectively). Gp53 was used for alignment of the models and shown in red. Models are shown in surface representation, arrows indicate displacement of domains gp7_III_ and gp7_II_.

**Extended Data Figure 6.**
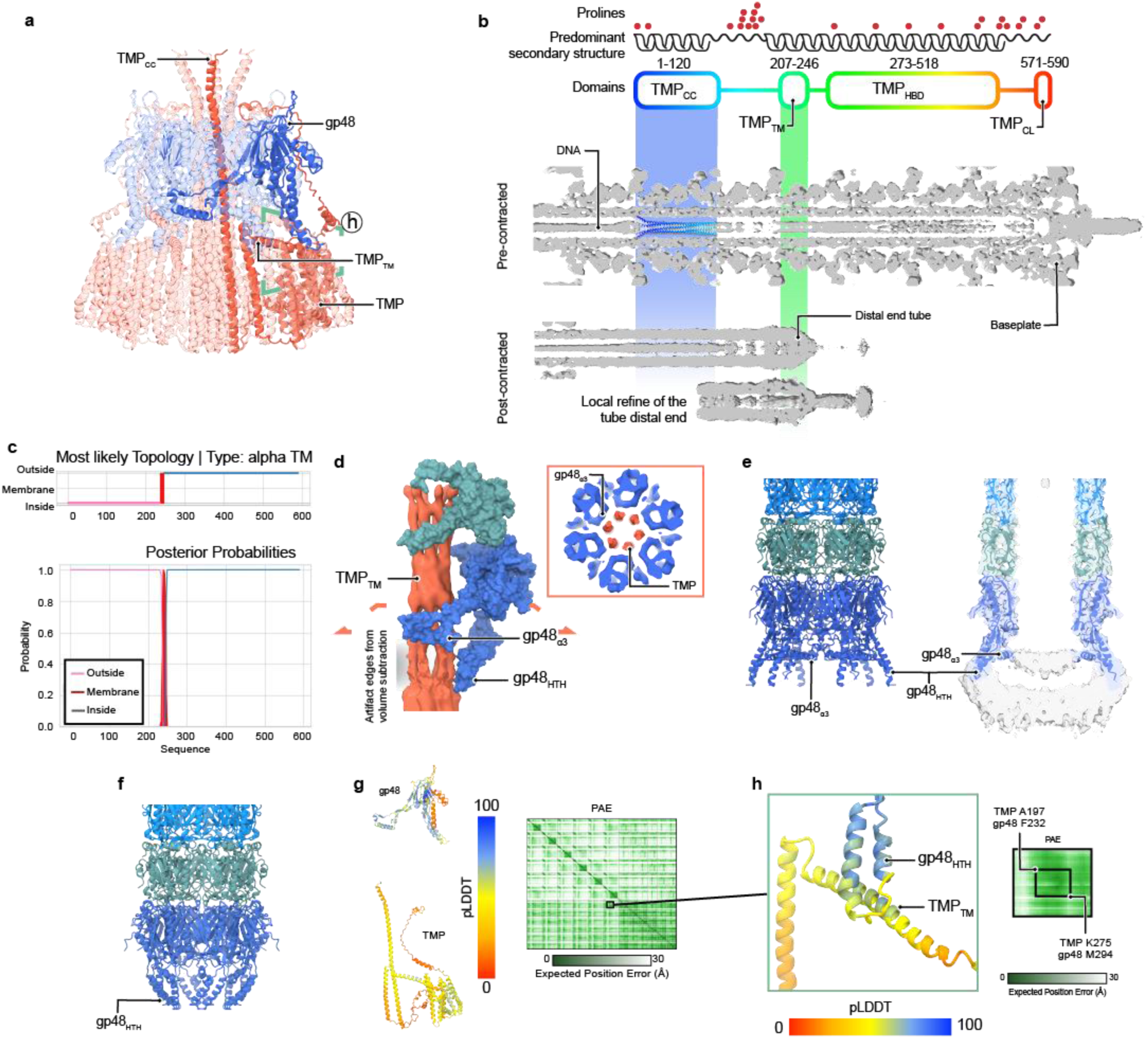
Tape measure protein and distal end tube structures. **a**, An AlphaFold2 prediction of the gp48_6_-TMP_6_ complex, prediction confidence ipTM= 0.625, highlighting one protomer of gp48 and TMP. **b**, Estimated arrangement of the TMP in the pre-and post-contracted states of the virion. The protein sequence shows a proline reach region between the TMP_CC_ and the TMP_TM_. The TMP_CC_ prediction from panel a is fitted in the bundle density for size reference. The precise location of the different domains could vary based on the level of packaging of the TMP inside the tube and is only shown for reference, however the TMP_TM_ location in the post-contracted tail can be approximated to the distal end tube. The distal end tube of the post-contracted state is shown from the full tail reconstruction and its local refinement. Reconstructions are at scale and aligned at the genome terminus. **c,** Surface representation of one gp48 and gp54 protomers and extracted cryo-EM reconstruction of the TMP in the post-contracted state before genome ejection. The inlet shows a transversal section of the cryo-EM reconstruction at the height of the gp48_α3_. **d,** Alternative state of the distal end tube after genome ejection, showing the flared gp48 in ribbon representation (*left*) and a section of the model fitted in the cryo-EM map at low threshold to show the unmodelled density at the end of the tube (*right*). **e**, Alternative state of the distal tube acter genome ejection where the gp48_a3_ could not be modelled. **f**, Focused view of the predicted interaction between the gp48_HTH_ and the TMP_TM_. The PAE matrix corresponds to the squared region on panel b. **g**, One chain of gp48 and one chain of TMP from the prediction shown in panel a, coloured by pLDDT values and the PAE matrix for the full complex prediction. **h,** Transmembrane prediction graph from DeepTMHMM of the TMP.

**Extended Data Figure 7.**
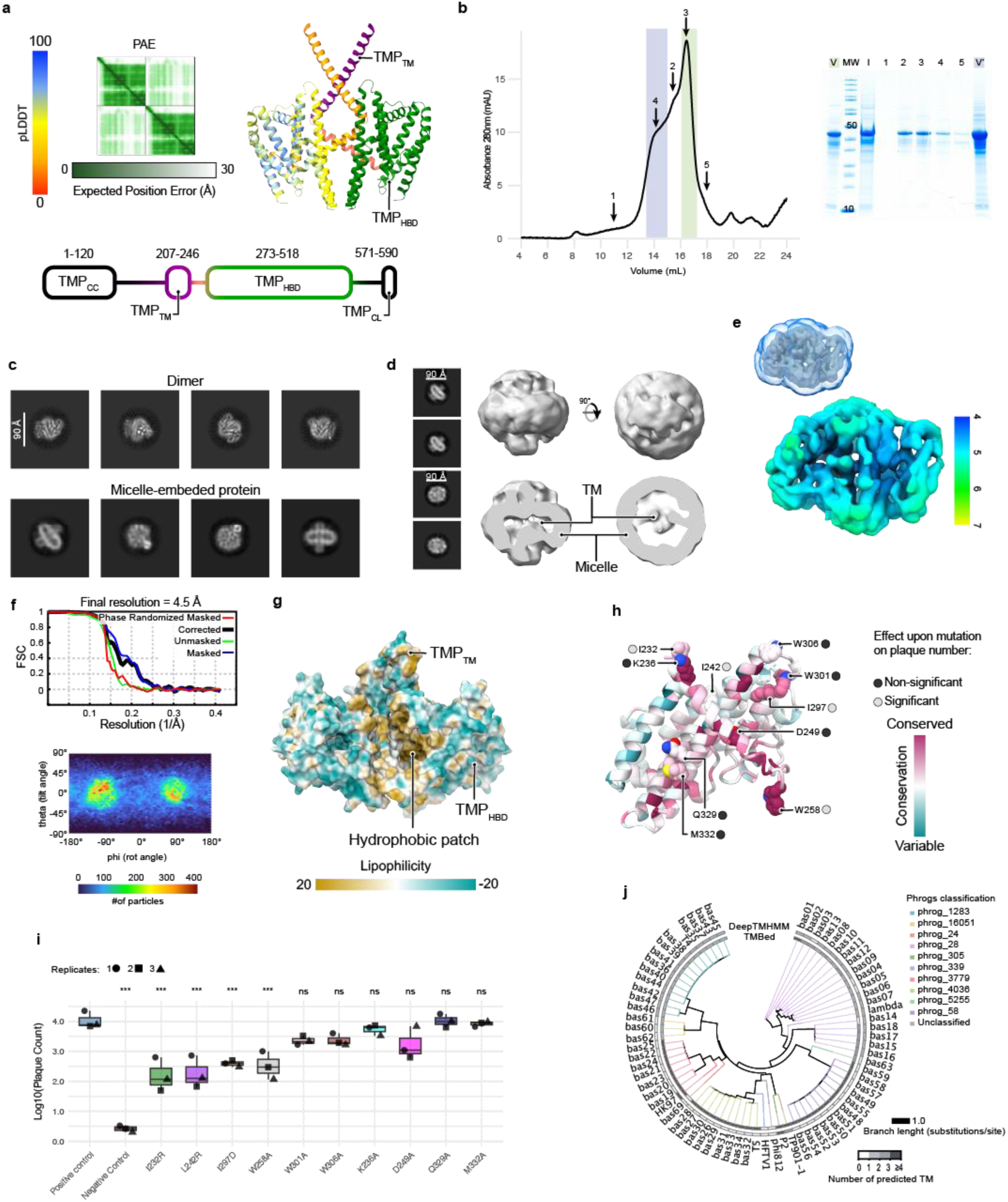
Construct of TMP_HBD_ and TMP_TM_ purification and structure analysis. a, Predicted TMP^S^ dimer and domain distribution. Left protomer coloured based on pLDDT and right protomer coloured based on domain distribution. **b,** Size-exclusion chromatography profile of purified TMP**^S^**and corresponding complete SDS-PAGE gel. Molecular weight (MW), input sample for the chromatography (I) and fractions (1 to 5) are also shown. V and V’ stand for the concentrated fractions used for vitrification as coloured in the elution profile. **c**, Representative 2D classes obtained from the picked particles that led to the dimer and representative 2D classes where a protein in a micelle can be observed. **d,** Ab-initio reconstruction of the 2D classes shown, arguably representing a micelle-embedded protein with transmembrane (TM) segments. **e,** Resolution and FSC curve of the cryo-EM reconstruction of the TMP^S^_2_. **f,** View distribution of reconstruction shown in panel e. **g,** Surface representation of the atomic model in panel c coloured based in lipophilicity **h,** One protomer of the TMP^S^_2_ coloured based on residue conservation, side chain of mutated residues are shown for reference. **i,** Plaque number obtained from the recombination of the phage T4 encoding the amber *g29* with homology vectors encoding the indicated mutations. Data are the mean of three replicates. **j,** Transmembrane prediction of the TMP proteins of the labelled phages using DeepTMHMM and TMBed. Number of predicted TM segments is coloured as shown and branches are coloured based on phrogs classification of the TMP protein.

**Extended Data Figure 8.**
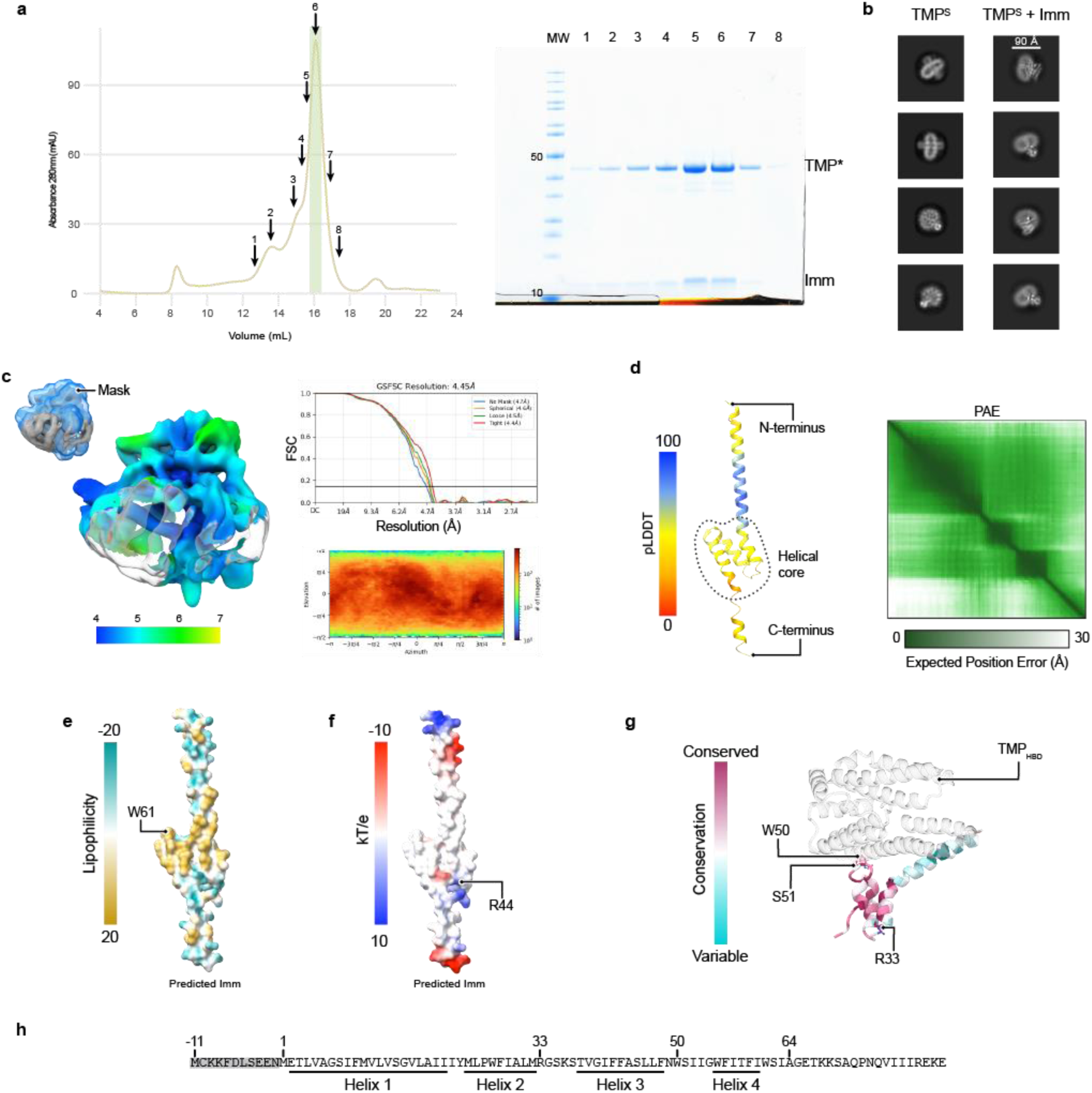
Purification of the TMP^S^-Imm complex and Imm characteristics. **a,** Size-exclusion chromatography profile of purified TMP**^S^**-Imm and corresponding complete SDS-PAGE gel. Molecular weight (MW) and fractions (1 to 8) are also shown. Concentrated fractions used for vitrification are coloured in the elution profile. **b,** Comparison between the 2D classes observed in the dataset of the TMPS and the TMPS-Imm. **c,** Resolution, FSC curve, and view distribution of the cryo-EM reconstruction. **d,** Ribbon representation of AlphaFold3 prediction of the 94-residue Imm protein used in cryo-EM data collection in this study coloured by pLDDT and PAE matrix shown. **e,** Same predicted model as in panel d with surface representation coloured based on lipophilicity. **f**, Same predicted model as in panel d with surface representation coloured based on electrostatic charge. **g,** Conservation profile of Imm in the atomic model of TMP-Imm refined in this study, showing the mutated residues and their effect on EOP of phage T4. **h,** Amino acid sequence of Imm, with the alternative early start codon transcribed in the grey area and numbered relevant residues in the present study.

**Extended Data Figure 9.**
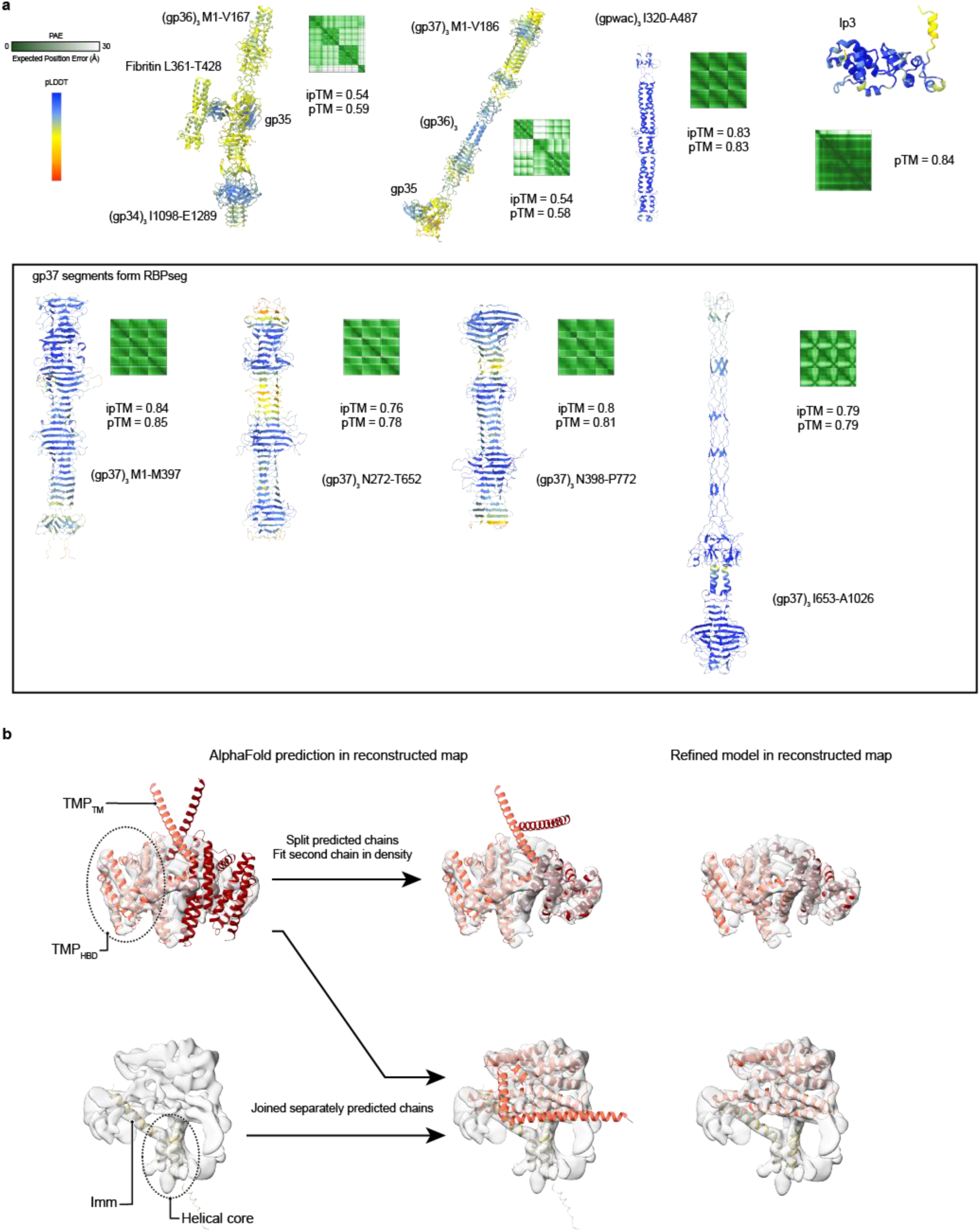
Additional information on predicted structures. **a,** AlphaFold predictions used in this study not shown previously, including the segments obtained from RBPseg before joining. **b,** Summary of the model building for the low-resolution maps of the TMP^S^_2_ and TMP-Imm, showing initial prediction fitting into the map compared to the final refined model. The pLDDT and PAE of the predictions are shown in Extended Data Figures 9 and 10 respectively.

**Supplementary Table 1.** Summary of the mass spectrometry results. This file contains the protein list for the proteins identified from the mass spectrometry data, together with the peptides and summary information of the mass spectrometry results. The full data can be found in the ProteimeXchange Consortium via the PRIDE partner repository with the dataset identifier PXD072569.

**Supplementary Table 2.** Cryo-EM data collection, refinement and validation statistics.

**Supplementary Table 3.** Plaque counting of g29 recombination and its mutants. This file contains the data of the counted plaques for the recombination experiment shown in Extended Data Fig. 9h,i.

**Supplementary Table 4.** Plaque counting of superinfection exclusion performance by Imm and its mutants. This file contains the data of the counted plaques for the measurements of superinfection exclusion infection efficiency of the Imm an its mutants shown in Fig. 5f.

**Supplementary Movie 1.** Schematic model of T4 early infection stages and proposed mechanism of genome delivery.

## Notes

### Competing Interest Statement

The authors have declared no competing interest.

### Summary of Updates

Minor corrections in text (grammar and spelling) and figures. The video summarising the phage T4 early infection is also included in this submission.

