## Extended data for "‘Genome delivery of a contractile tailed phage and its superinfection exclusion mechanism’"

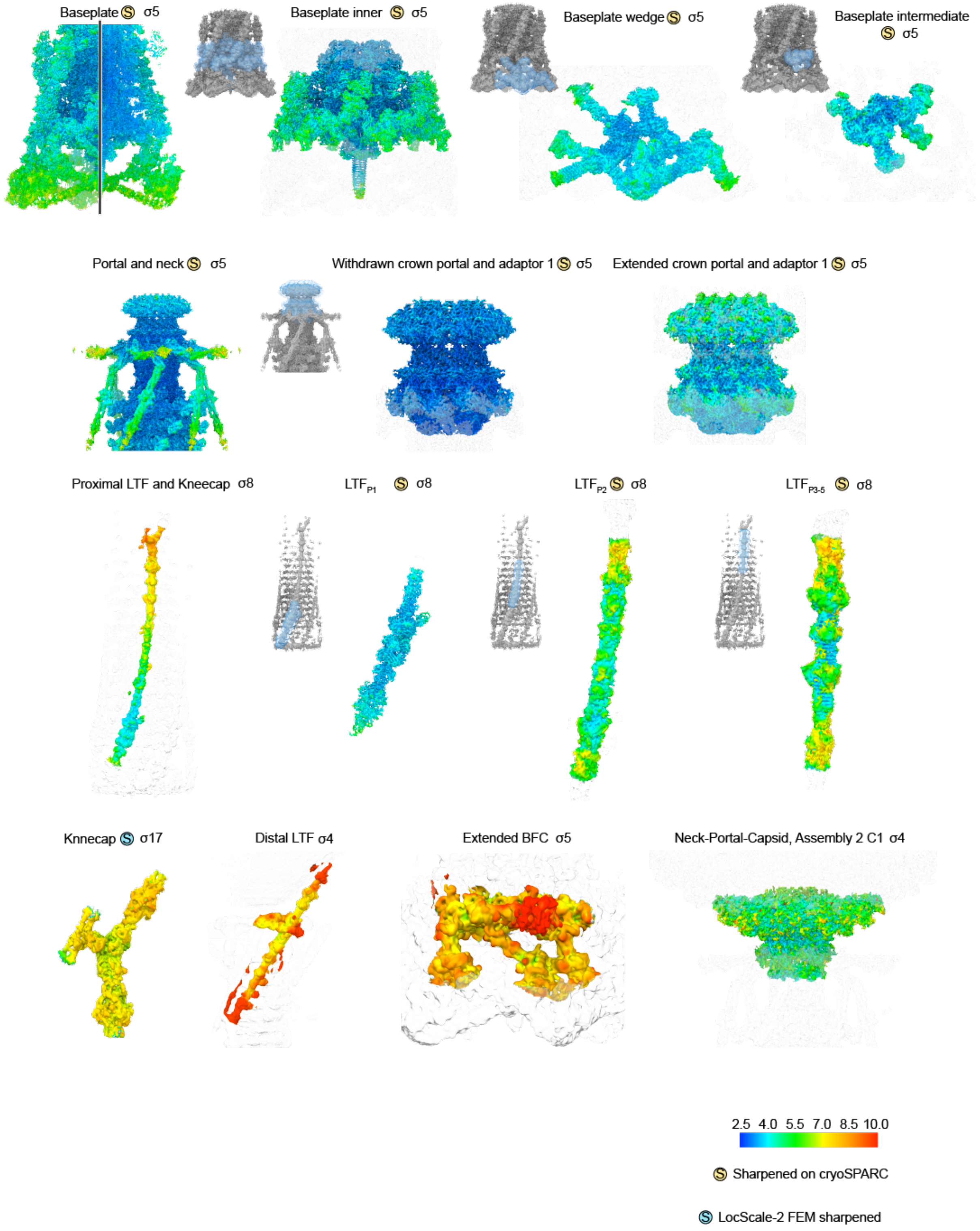


**Extended Data Figure 1. Local resolution estimation of the reconstructions used for model building of the pre-contracted virion.**

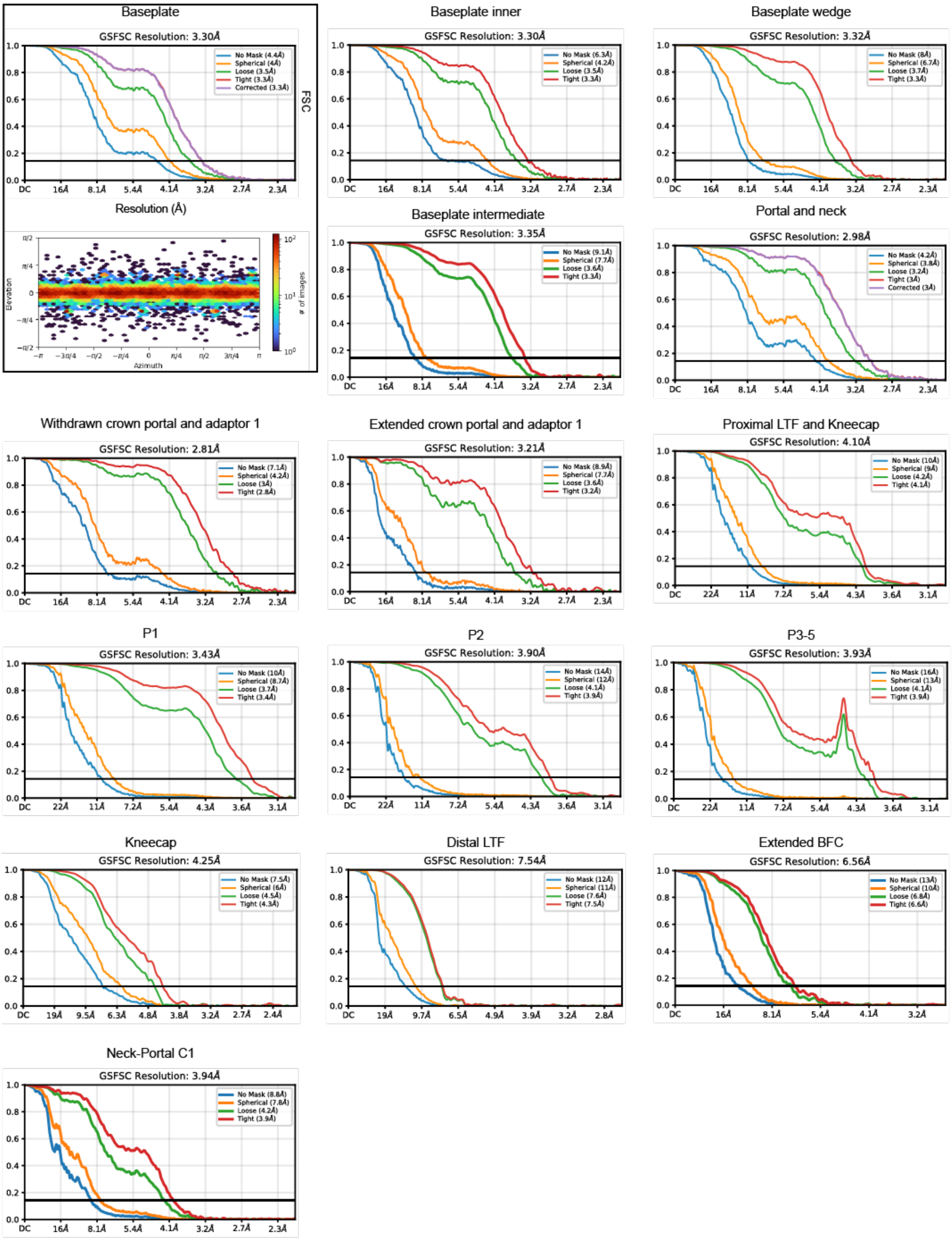


**Extended Data Figure 2. FSC curves of the reconstructions of the pre-contracted virion.**

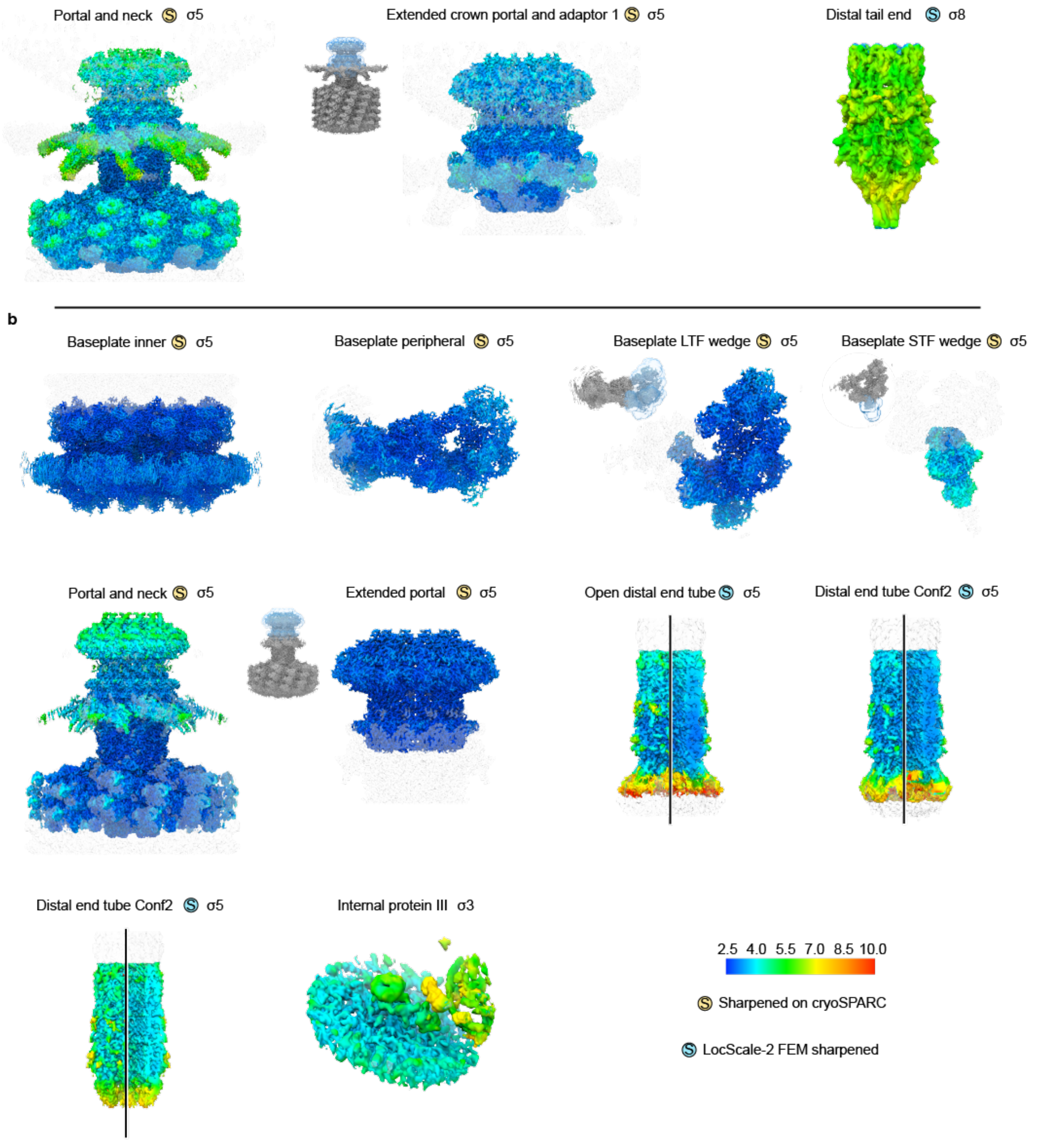


**Extended Data Figure 3. Local resolution estimation of the reconstructions used for model building of the post-contracted virion and post-ejected genome virion. a,** Reconstructions of the post-contracted virion. **b,** Reconstructions of the post-ejected genome virion. It is indicated when the reconstructions have been sharpened either with cryoSPARC or LocScale.

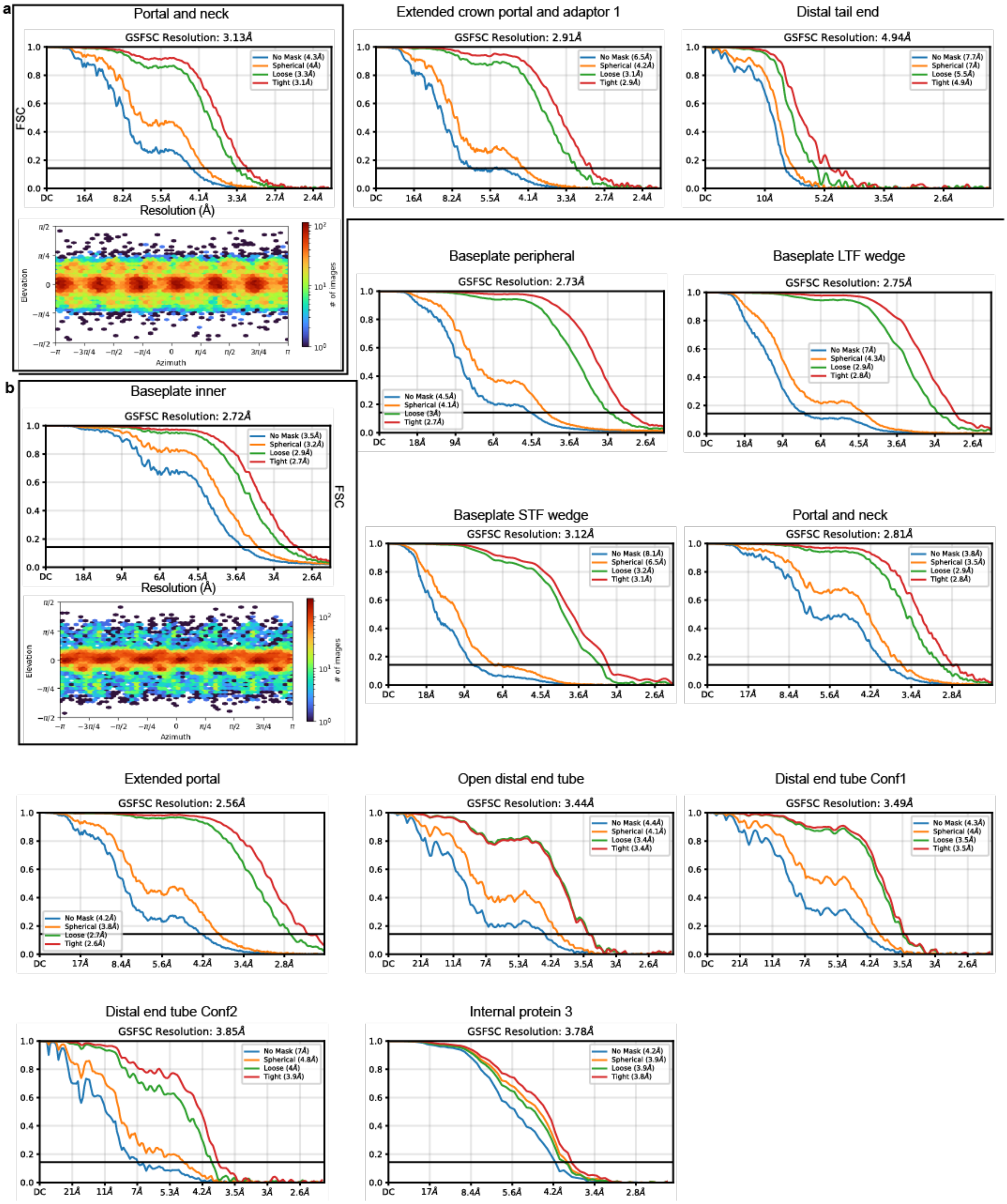


**Extended Data Figure 4. FSC curves of the reconstructions of the post-contracted virion and post-ejected genome virion. a,** FSC curves of the post-contracted virion reconstructions. **b,** FSC curves of the post-ejected genome virion reconstructions.

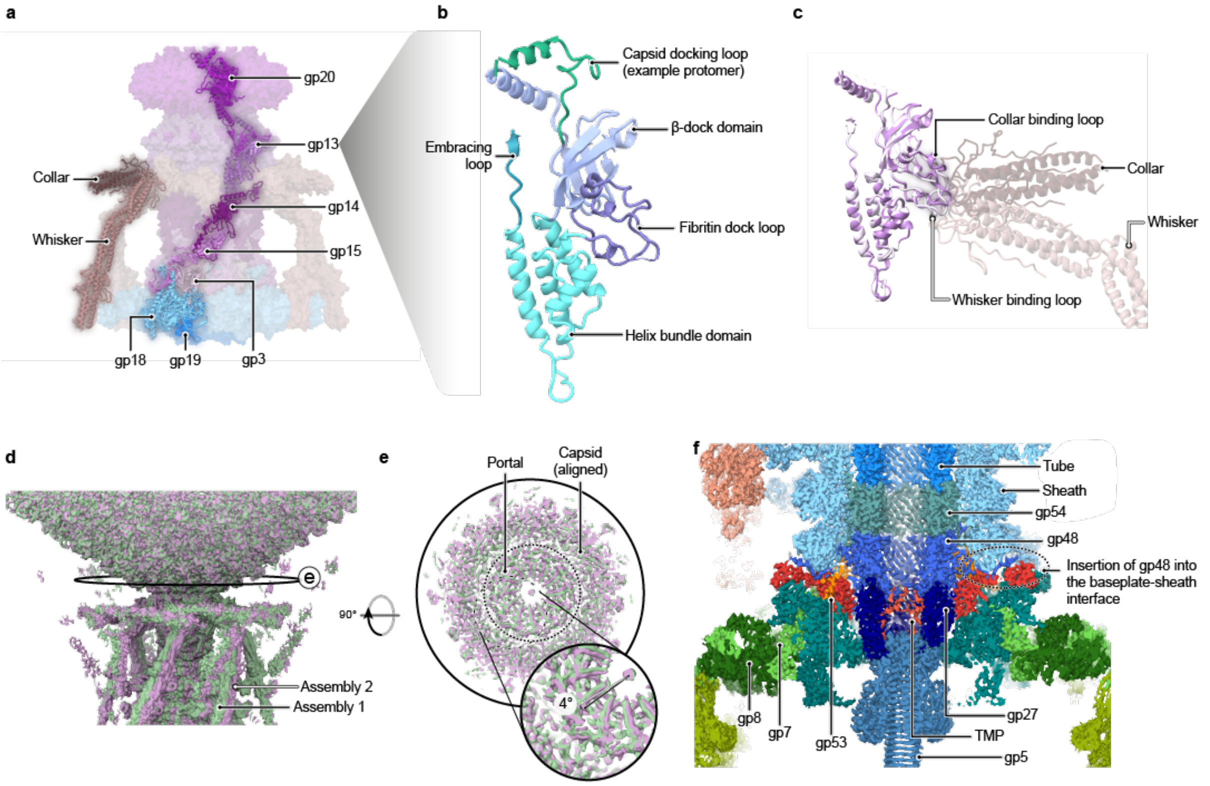


**Extended Data Figure 5. Phage T4 virion architecture. a,** Organisation of the neck–portal complex of phage T4. **b,** Structure of a representative monomer of gp13, the neck adaptor. The domains are subdivided based on the gp11 of phage SU10[40] and gp81 of phage GP4[41], which were matched by FoldSeek with E-values of 6.11e-2 and 1.03e-1 respectively. **c,** Comparison of one gp13 monomer binding the collar and another binding the whisker. **d,** Alignment of the cryo-EM reconstruction of the two tail-capsid assemblies by alignment of the capsid density. **e,** Clipped top-view of the same alignment as in panel d. **f,** Sliced cryo-EM reconstruction of the inner and intermediate baseplate, and distal tail end.

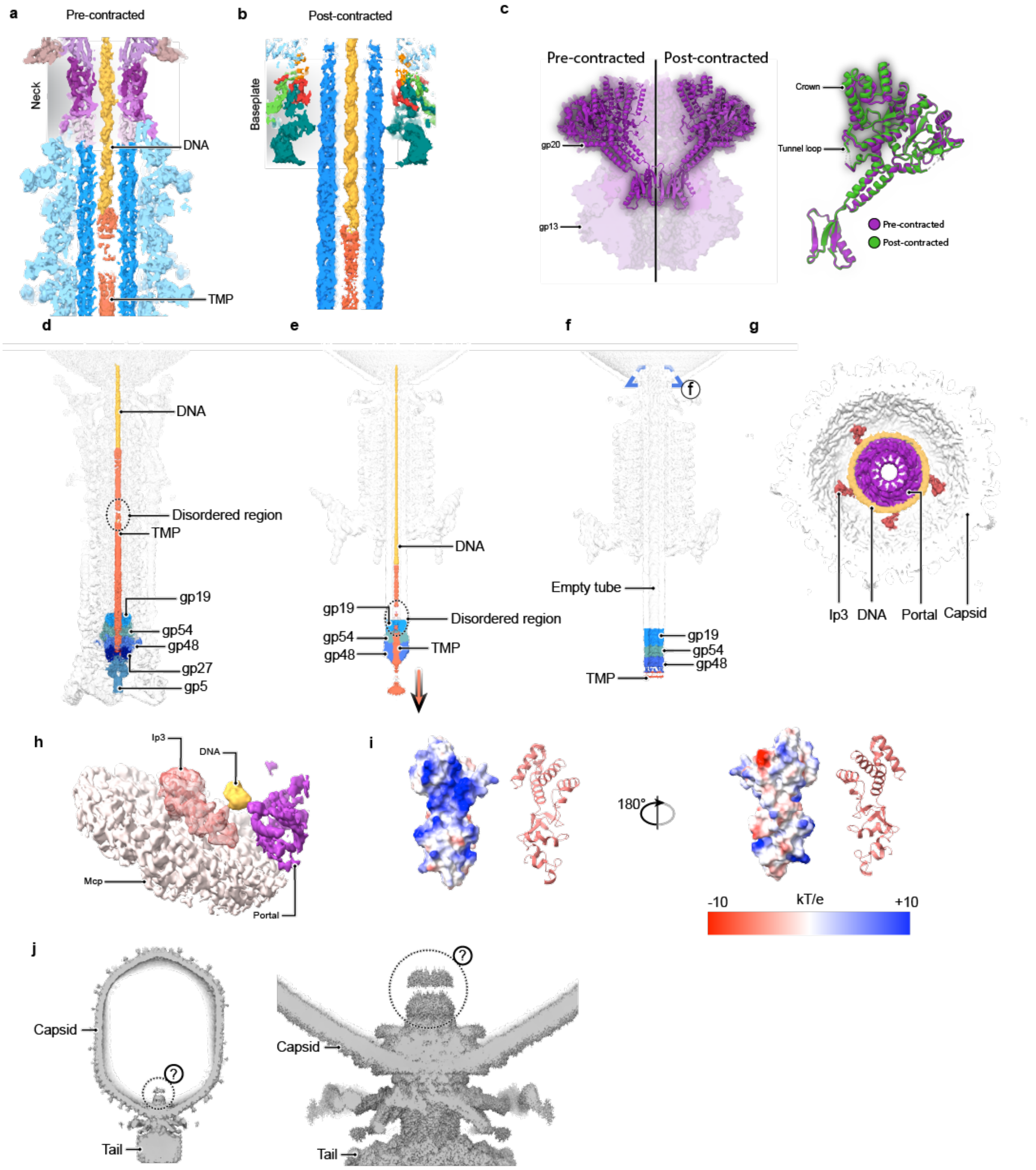


**Extended Data Figure 6. Overview of genome delivery process**. **a,** Sliced cryo-EM map of the pre-contracted virion at the tube proximal end, where the genome terminus binds to the TMP. **b,** Sliced cryo-EM map of the post-contracted virion at the level of the baseplate, where the genome terminus is retained after contraction. **c,** Comparison between the pre- and post-contracted conformations of the portal protein gp20 and alignment of both portal states. **d,** Sliced cryo-EM map of complete tail and neck of phage T4 in pre-contracted state, where DNA, TMP, baseplate proximal tube, and spike are coloured. **e,** Post-contracted virion state represented as in panel a. Arrow indicates ejection of the TMP. **f**, Post-ejected genome virion state in presence of nanodiscs, represented as in panel e. **g**, View of the portal in the post-ejected genome capsid, showing the density of DNA around the portal and the Ip3 copies (Ip3, internal protein III). **h,** Cryo-EM reconstruction of the local refinement of the Ip3, refined AlphaFold3 prediction fitted in the density. **i,** Surface representation of the Ip3 coloured based on electrostatic potential. **j,** Reconstruction of the capsid without symmetry imposed (*left*) and neck-portal with six-fold symmetry imposed (*right*) at low threshold to show the unresolved density above the portal.

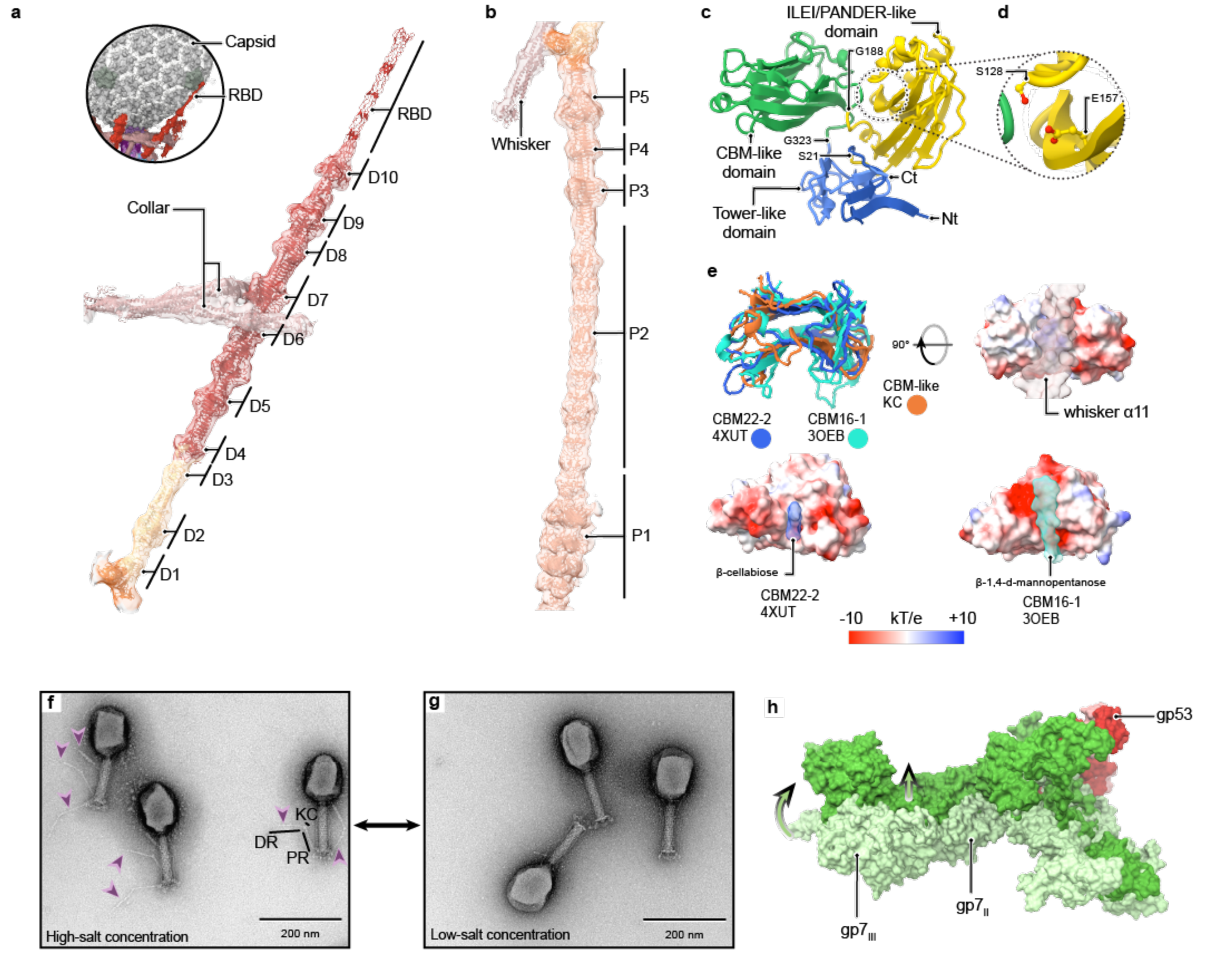


**Extended Data Figure 7. Fibre architecture and extension**. **a**, Distal rod, kneecap and collar fibritins cryo-EM map and the refined model fitted. Two collar fibritins clamp $(gp37)_3$ in between domains 6 and 7. The RBD could not be resolved for all the distal rods as result of the different bends due to the capsid, but one RBD was bound to the capsid and present on the capsid refinement (*top left*) (RBD, receptor binding domain) **b**, Proximal rod, kneecap and whisker fibritin cryo-EM map and the refined model fitted. **c**, Domain distribution of the kneecap protein. **d**, Detailed view of the conserved hydrogen bound in the ILEI/PANDER-like domain[46]. **e**, Aligned structure of the CBM-like domain of the kneecap to the CBM22-2 and CBM16-1. Each structure is also surface represented facing the groove, where the ligand is found (the whisker α11 for the kneecap) and coloured based on electrostatic potential. The TM-score for the CBM22-2 is 0.712 and RMSD=2.64 Å for 115 residues, and for CBM16-1 is 0.729 and RMSD=2.16 Å for 109 residues. **f**, TEM image of negative stained phage T4 sample in high-salt buffer. Purple arrows indicate extended LTF. **g**, Same sample as imaged in panel g, but after buffer exchange to low-salt buffer. **h**, Gp7 conformational change from the pre-contracted baseplate and the post-contracted state (light and darker colours respectively). Gp53 was used for alignment of the models and shown in red. Models are shown in surface representation, arrows indicate displacement of domains $gp7_{III}$ and $gp7_{II}$.

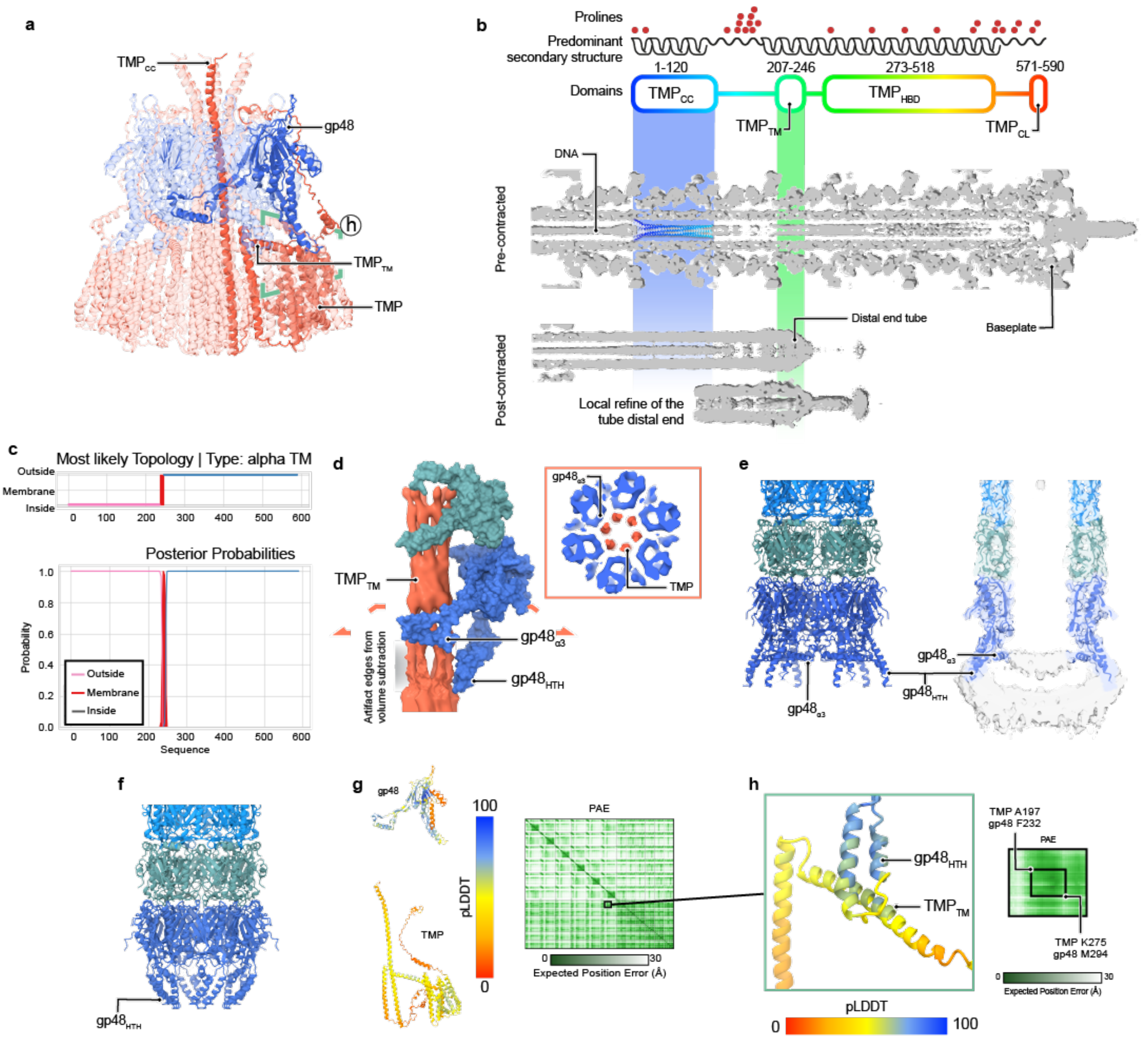


**Extended Data Figure 8. Tape measure protein and distal end tube structures**. **a**, An AlphaFold2 prediction of the $gp48_6$-$TMP_6$ complex, prediction confidence ipTM= 0.625, highlighting one protomer of gp48 and TMP. **b**, Estimated arrangement of the TMP in the pre- and post-contracted states of the virion. The protein sequence shows a proline reach region between the $TMP_{CC}$ and the $TMP_{TM}$. The $TMP_{CC}$ prediction from panel a is fitted in the bundle density for size reference. The precise location of the different domains could vary based on the level of packaging of the TMP inside the tube and is only shown for reference, however the $TMP_{TM}$ location in the post-contracted tail can be approximated to the distal end tube. The distal end tube of the post-contracted state is shown from the full tail reconstruction and its local refinement. Reconstructions are at scale and aligned at the genome terminus. **c,** Surface representation of one gp48 and gp54 protomers and extracted cryo-EM reconstruction of the TMP in the post-contracted state before genome ejection. The inlet shows a transversal section of the cryo-EM reconstruction at the height of the gp48$_{\alpha3}$. **d,** Alternative state of the distal end tube after genome ejection, showing the flared gp48 in ribbon representation (*left*) and a section of the model fitted in the cryo-EM map at low threshold to show the unmodelled density at the end of the tube (*right*). **e**, Alternative state of the distal tube acter genome ejection where the gp48$_{a3}$ could not be modelled. **f**, Focused view of the predicted interaction between the gp48$_{HTH}$ and the TMP$_{TM}$. The PAE matrix corresponds to the squared region on panel b. **g**, One chain of gp48 and one chain of TMP from the prediction shown in panel a, coloured by pLDDT values and the PAE matrix for the full complex prediction. **h,** Transmembrane prediction graph from DeepTMHMM of the TMP.

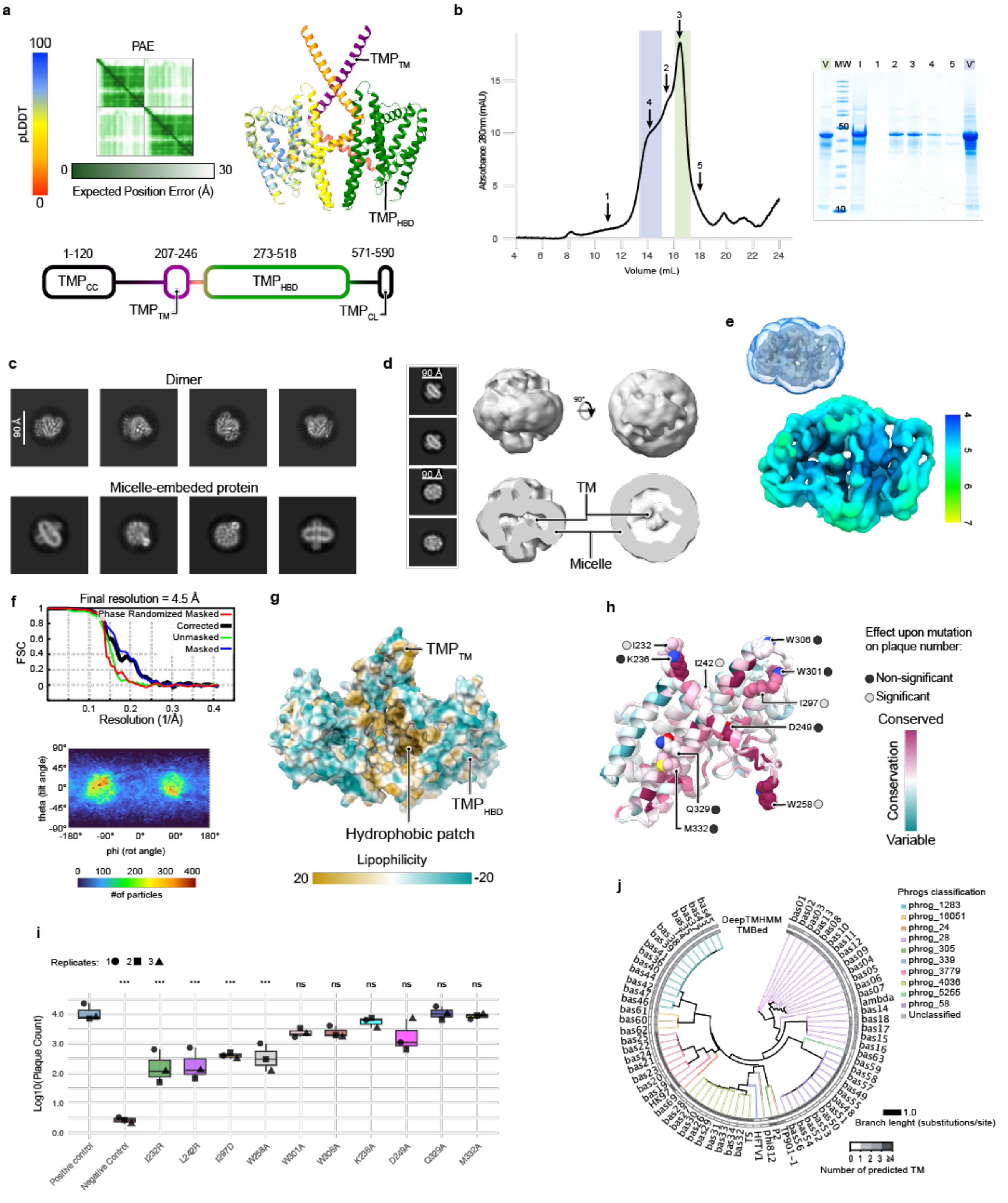


**Extended Data Figure 9. Construct of TMP$_{HBD}$ and TMP$_{TM}$ purification and structure analysis. a,** Predicted TMP$^S$ dimer and domain distribution. Left protomer coloured based on pLDDT and right protomer coloured based on domain distribution. **b,** Size-exclusion chromatography profile of purified TMP$^S$ and corresponding complete SDS-PAGE gel. Molecular weight (MW), input sample for the chromatography (I) and fractions (1 to 5) are also shown. V and V' stand for the concentrated fractions used for vitrification as coloured in the elution profile. **c**, Representative 2D classes obtained from the picked particles that led to the dimer and representative 2D classes where a protein in a micelle can be observed. **d,** Ab-initio reconstruction of the 2D classes shown, arguably representing a micelle-embedded protein with transmembrane (TM) segments. **e,** Resolution and FSC curve of the cryo-EM reconstruction of the $TMP^{S}_{2}$. **f,** View distribution of reconstruction shown in panel e. **g,** Surface representation of the atomic model in panel c coloured based in lipophilicity **h,** One protomer of the $TMP^{S}_{2}$ coloured based on residue conservation, side chain of mutated residues are shown for reference. **i,** Plaque number obtained from the recombination of the phage T4 encoding the amber *g29* with homology vectors encoding the indicated mutations. Data are the mean of three replicates. **j,** Transmembrane prediction of the TMP proteins of the labelled phages using DeepTMHMM and TMBed. Number of predicted TM segments is coloured as shown and branches are coloured based on phrogs classification of the TMP protein.

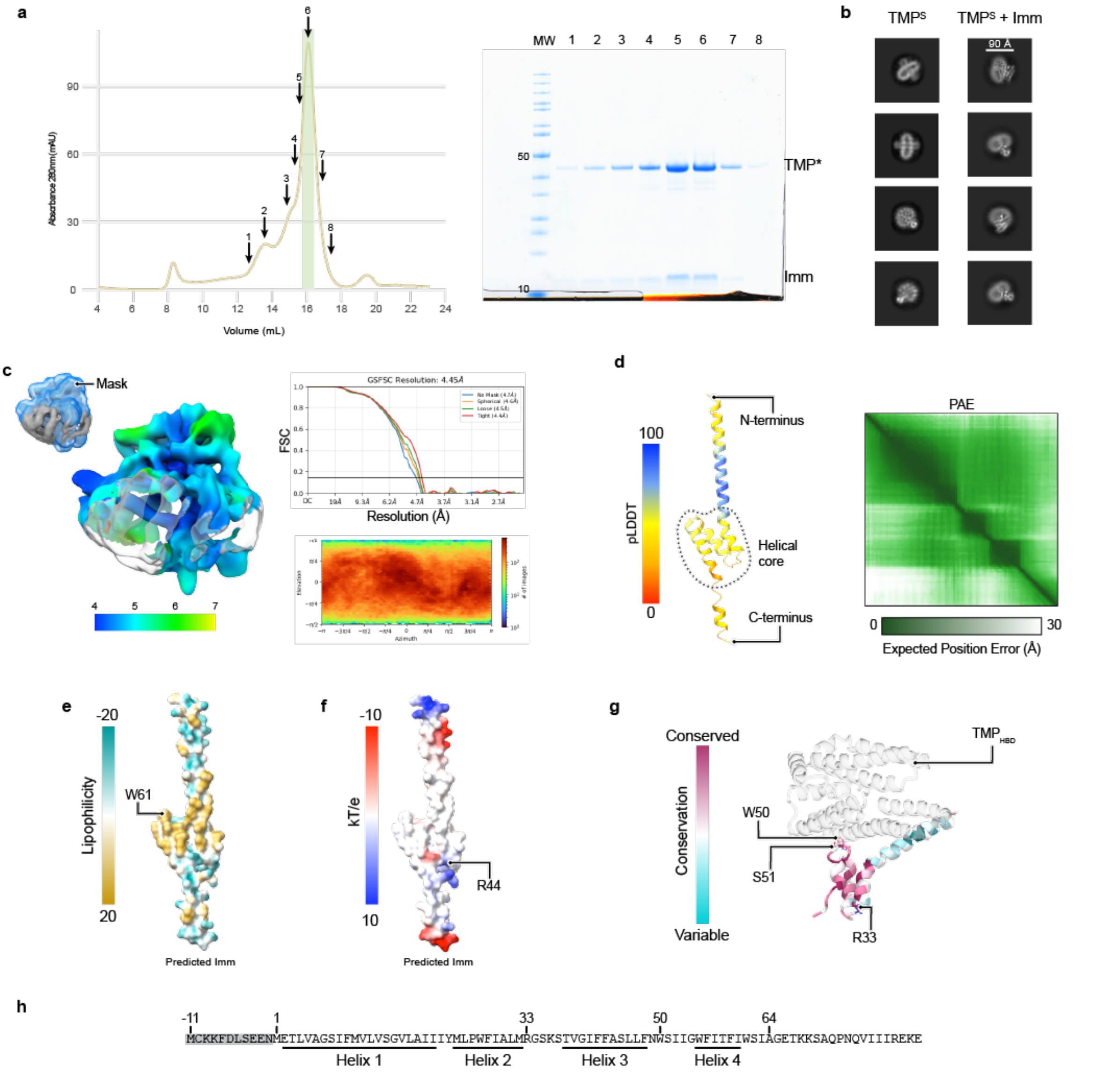


**Extended Data Figure 10. Purification of the TMP$^S$-Imm complex and Imm characteristics**. **a,** Size-exclusion chromatography profile of purified TMP$^S$-Imm and corresponding complete SDS-PAGE gel. Molecular weight (MW) and fractions (1 to 8) are also shown. Concentrated fractions used for vitrification are coloured in the elution profile. **b,** Comparison between the 2D classes observed in the dataset of the TMPS and the TMPS-Imm. **c,** Resolution, FSC curve, and view distribution of the cryo-EM reconstruction. **d,** Ribbon representation of AlphaFold3 prediction of the 94-residue Imm protein used in cryo-EM data collection in this study coloured by pLDDT and PAE matrix shown. **e,** Same predicted model as in panel d with surface representation coloured based on lipophilicity. **f**, Same predicted model as in panel d with surface representation coloured based on electrostatic charge. **g,** Conservation profile of Imm in the atomic model of TMP-Imm refined in this study, showing the mutated residues and their effect on EOP of phage T4. **h,** Amino acid sequence of Imm, with the alternative early start codon transcribed in the grey area and numbered relevant residues in the present study.

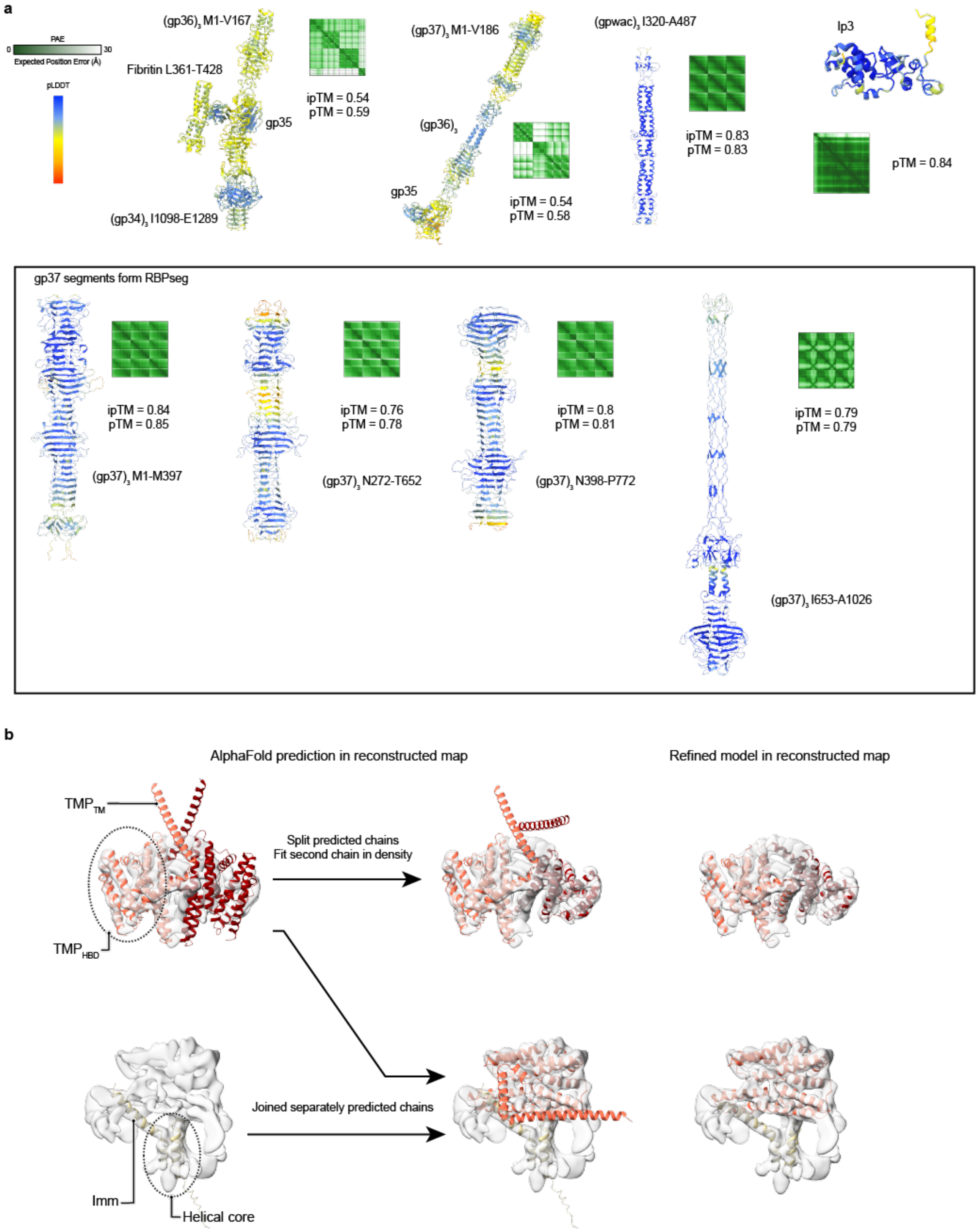


**Extended Data Figure 11. Additional information on predicted structures. a,** AlphaFold predictions used in this study not shown previously, including the segments obtained from RBPseg before joining. **b,** Summary of the model building for the low-resolution maps of the $TMP^S_2$ and TMP-Imm, showing initial prediction fitting into the map compared to the final refined model. The pLDDT and PAE of the predictions are shown in Extended Data Figures 9 and 10 respectively.

**Supplementary Table 1.** Summary of the mass spectrometry results. This file contains the protein list for the proteins identified from the mass spectrometry data, together with the peptides and summary information of the mass spectrometry results. The full data can be found in the ProteimeXchange Consortium via the PRIDE partner repository with the dataset identifier PXD072569.

**Supplementary Table 2.** Cryo-EM data collection, refinement and validation statistics.

**Supplementary Table 3.** Plaque counting of g29 recombination and its mutants. This file contains the data of the counted plaques for the recombination experiment shown in Extended Data Fig. 9h,i.

**Supplementary Table 4.** Plaque counting of superinfection exclusion performance by Imm and its mutants. This file contains the data of the counted plaques for the measurements of superinfection exclusion infection efficiency of the Imm an its mutants shown in Fig. 5f.

**Supplementary Movie 1.** Schematic model of T4 early infection stages and proposed mechanism of genome delivery.
